# Human back-tracking behaviour is associated with suppression of default-mode region activity and enhanced dorsal anterior cingulate activity

**DOI:** 10.1101/362319

**Authors:** Amir-Homayoun Javadi, Eva Zita Patai, Eugenia Marin-Garcia, Aaron Margois, Heng-Ru M. Tan, Dharshan Kumaran, Marko Nardini, Will Penny, Emrah Duzel, Peter Dayan, Hugo J. Spiers

## Abstract

Central to the concept of the ‘cognitive map’ is that it confers behavioural flexibility, allowing animals to take efficient detours, exploit shortcuts and realise the need to back-track rather than persevere on a poorly chosen route. The neural underpinnings of such naturalistic and flexible behaviour remain unclear. During fMRI we tested human subjects on their ability to navigate to a set of goal locations in a virtual desert island riven by lava, which occasionally shifted to block selected paths (necessitating detours) or receded to open new paths (affording shortcuts). We found that during self-initiated back-tracking, activity increased in frontal regions and the dorsal anterior cingulate cortex, while activity in regions associated with the core default-mode network was suppressed. Detours activated a network of frontal regions compared to shortcuts. Activity in right dorsolateral prefrontal cortex specifically increased when participants encountered new plausible shortcuts but which in fact added to the path (false shortcuts). These results help inform current models as to how the brain supports navigation and planning in dynamic environments.

**Significance Statement:** Adaptation to change is important for survival. Although real-world spatial environments are prone to continual change, little is known about how the brain supports navigation in dynamic environments where flexible adjustments to route plans are needed. Here, we used fMRI to examine the brain activity elicited when humans took forced detours, identified shortcuts and spontaneously back-tracked along their recent path. Both externally and internally generated changes in the route activated the fronto-parietal attention network, whereas only internally generated changes generated increased activity in the dorsal anterior cingulate cortex with a concomitant disengagement in regions associated with the default-mode network. The results provide new insights into how the brain plans and re-plans in the face of a changing environment.

## Introduction

A challenge all motile animals face is adapting to changes in an environment so that they can efficiently return to safety or find food. Adaptations include identifying novel shortcuts and minimizing the lengths of imposed detours. Tolman (1948) conceptualized this ability as arising from an internal ‘cognitive map’ (or, in control theoretic terms, an internal model) of the environment. Evidence from electrophysiological recordings in rodents and fMRI in humans has supported the view that hippocampus contains a cognitive map (Ekstrom, Spiers, Bohbot, & Rosenbaum, 2018; Epstein, Patai, Julian, & Spiers, 2017; O’Keefe & Nadel, 1978; Spiers & Barry, 2015). However, the evidentiary basis in humans and other animals of the navigational functions of the cognitive map in the face of dynamic environments is incomplete.

Early studies in rodents (Tolman & Honzik, 1930) along with more recent studies in rats and other mammals (Alvernhe, Save, & Poucet, 2011; Alvernhe, Van Cauter, Save, & Poucet, 2008; Chapuis, 1987; Chapuis, Durup, & Thinus-Blanc, 1987; Poucet, Thinus-Blanc, & Chapuis, 1983; Winocur, Moscovitch, Rosenbaum, & Sekeres, 2010) have helped characterise flexible navigation behaviour when the environmental layout changes. However, there has been relatively limited investigation of the evoked neural responses at the moment of encountering changes in the environment. By contrast, a number of functional neuroimaging studies in humans have studied the evoked responses to detours (Howard et al., 2014; Iaria, Fox, Chen, Petrides, & Barton, 2008; Maguire et al., 1998; Rauchs et al., 2008; Rosenbaum, Ziegler, Winocur, Grady, & Moscovitch, 2004; Simon & Daw, 2011; Spiers & Maguire, 2006; Xu, Evensmoen, Lehn, Pintzka, & Håberg, 2010). Rather than revealing hippocampal activity, these studies have consistently reported increased prefrontal activity. These studies report: i) increased activity in right lateral prefrontal regions when detecting changes in the environment, ii) activity in frontopolar cortex when re-planning and setting sub-goals, and iii) superior prefrontal cortical activity when processing conflict between route options (Spiers & Gilbert, 2015). Such responses are consistent with the view that the prefrontal cortex (PFC) supports flexible behaviour in response to changing affordances in the environment (Shallice, 1982; Spiers, 2008).

While some functional magnetic resonance imaging studies have examined shortcuts and detours in changing environments (Simon & Daw, 2011; Yoshida & Ishii, 2006) the paradigms deployed were not optimized to disentangle the neural response to detours and shortcuts, thus to date we lack evidence for how neural systems adjust at shortcuts and how they compare to detours. Since both detours and shortcuts change the path to the goal it is possible that both events elicit a similar neural response. Alternatively, the potential to simulate a new alternative path at detours might evoke increased activity in frontal regions associated with planning. Similarly, the process of determining that a newly encountered path is not in fact a useful route (a false shortcut), and needs to be suppressed, may draw on greater neural resources than when new path is identified as viable. Finally, some re-planning in the real-world is purely spontaneous, self-driven (Spiers and Maguire 2006) where an optimal action is to backtrack along the recent path to reach the goal. To date this behaviour has mainly been studied in ants (Wystrach, Schwarz, Baniel, & Cheng, 2013) and thus the neural correlates of such back-tracking currently remain elusive. To explore neural responses during back-tracking, detours and shorcuts we combined fMRI with a virtual reality (VR)-based environment (‘LavaWorld’) in which participants navigated a desert island containing hidden treasure with paths constrained by lava, which had the capacity to shift and open new paths (shortcuts) or close others off (detours).

## Methods

### Participants

Twenty-two subjects (mean age: 21.8 ± 2.3 years, range: 19-27; 14 female). Participants were administered a questionnaire regarding their navigation abilities/strategies (Santa Barbara Sense of Direction Scale; mean score = 4.9, range: 3.7-5.7). All participants had normal to corrected vision, reported no medical implant containing metal, no history of neurological or psychiatric condition, color blindness, and did not suffer from claustrophobia.

All participants gave written consent to participate to the study in accordance with the Birkbeck-UCL Centre for Neuroimaging ethics committee. Subjects were compensated with a minimum of £70 plus an additional £10 reward for good performance during the scan. One participant was excluded from the final sample because there was severe signal loss from the medial-temporal area in their functional scan.

### VR environment: Lavaworld

A virtual island maze environment was created using Vizard virtual reality software (© WorldViz). The maze was a grid network, consisting of ‘sand’ areas that were walkable, and ‘lava’ areas, which were unpassable and as such were like walls in a traditional maze. However, the whole maze layout was flat, so there was visibility into the distance over both sand and lava. This allowed participants to stay oriented in the maze throughout the task. Orientation cues were provided by four unique large objects in the distance. Movement was controlled by 4 buttons: left, right, forwards and backwards. Pressing left, right or backwards moved the participant to the grid square to the left, right or behind respectively (if there was no lava in the way), and rotated the view accordingly. Similarly, pressing forward moved the participant to the next square along. See Figure 1 for a participant viewpoint at one point in the maze. Participants were tested over two days, on day one they were trained on the maze, and on day two they were tested on the maze in the MRI scanner.

### Training

On the first day, participants were trained on the virtual maze (25 × 15 grid) to find goal locations. During this phase, all goal objects (20 in total, distributed across the maze) were visible at all times, and participants navigated from one to the next based on the currently displayed target object (displayed in the top-right corner of the screen). After one hour of training, subjects were given a test to establish how well they had learnt the object locations. On a blank grid, where only the lava was marked, participants had to place all the objects they remembered. They were given feedback from the experimenter, and if needed, prompts as to the missing objects. This memory-test was repeated twice more during the training, after 1.5 and 2 hours. At completion, for participants to return for the fMRI phase on the second day, they had to score at 100% accuracy in placing the objects.

**Figure 1:**
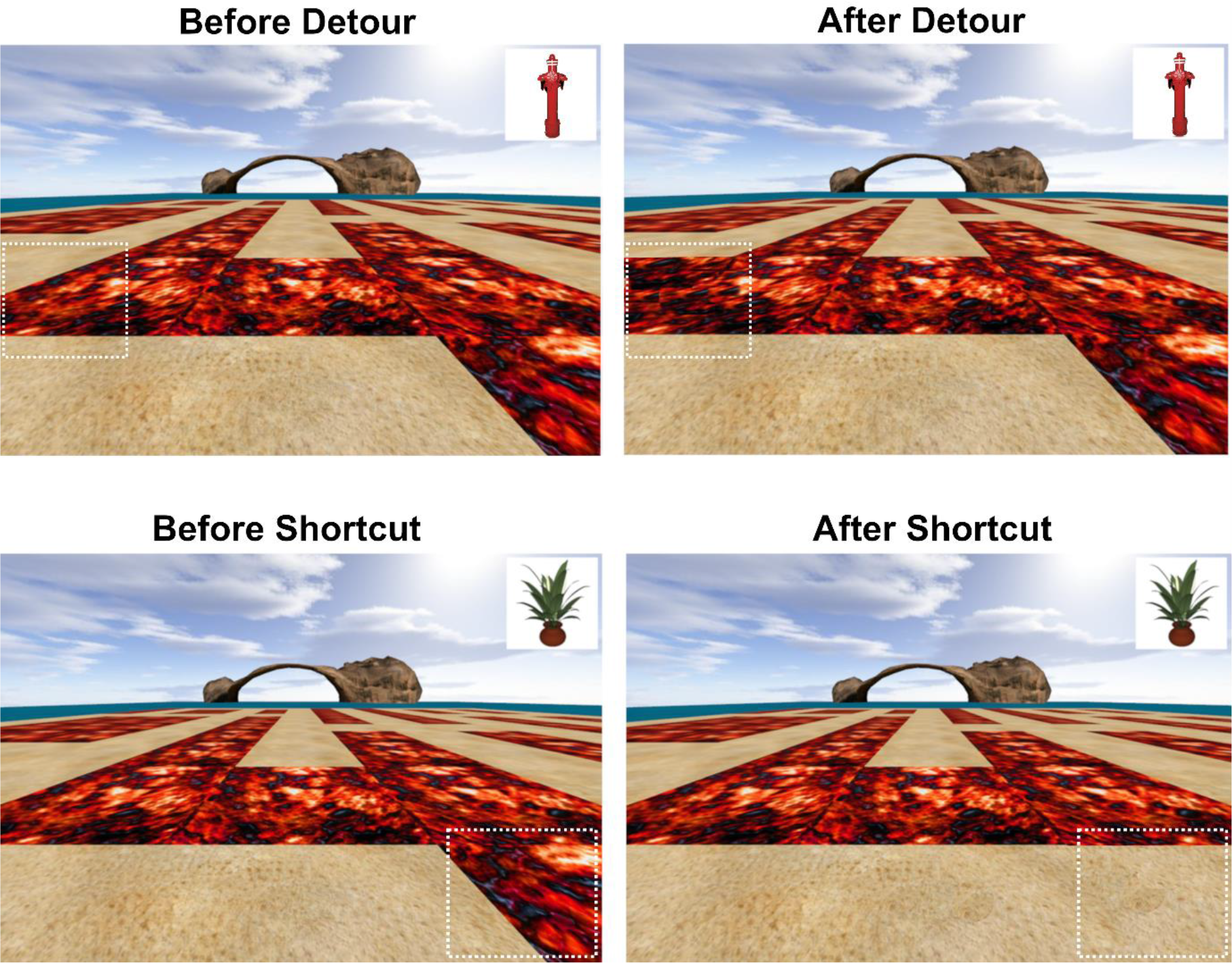
LavaWorld. Example view of test environment and current goal object (top right corner). A distal cue is visible (arch), and 3 others were located at the other cardinal directions. The sand represents the path that can be moved along, whereas the red ‘lava’ blocks in the path. During Training, objects were visible across the whole maze, and participants used the controls to move forward, left/right and backward to collect them, with an arrow guiding them towards the object (in the first of three rounds of training). During the Test phase, the objects were <u>not</u> visible and the environment could change, such that the lava shifted around to close an existing path (Detours, top row), or reveal new paths (Shortcuts or False Shortcuts, bottom row). White dotted boxes are to highlight the changes, and were not present during the experiment. See Figure 2 for more examples.

### Test & fMRI scan

On the test day, participants were given a brief refresher of the maze with the objects before beginning the test phase. Before scanning, participants were allowed to familiarize themselves with the scanner button pad, and the changes that would occur. This involved presenting them with a novel environment that had not been experienced on day one, and which had no objects, different distal cues and a different maze layout, to avoid any confounds or confusion with training and test mazes. Participants could then practice the task in this new environment, and accustom themselves to the controls (button pad with 4 active buttons: left, right, forward, and turn around) and to the appearance of changes to the lava.

While in the MRI scanner, participants performed the test phase of the experiment. A single trial in the test phase is defined as being informed which is the new goal object, and then finding the way to, and arriving at, it. During the test phase, two things were different from training: 1) target objects were not visible, so participants had to navigate between them based on their memories of the locations, and 2) the lava could move, blocking some paths and creating new paths. Participants were informed that this was a temporary change and that after reaching the goal the environment would revert to the baseline state. During each journey to an object, a single change event occurred in the lava layout. At the point of a change, the screen froze for 4 seconds to ensure that participants had an opportunity to detect the change and consider their path options. These changes could either be Detours (when a piece of lava was added to block the current path on the grid, thus forcing the participant to take an new longer route to their goal); Shortcuts (a piece of lava was removed and replaced with sand, allowing the participant to pick a shorter route); False Shortcuts (visually identical to Shortcuts, but choosing a route through them would increase the net distance to the goal because of the layout of the maze; False Shortcuts came in two classes: False Shortcuts Towards and False Shortcuts Away from the goal, depending on whether or not the False Shortcut seemed to lead in the general direction of the goal or if it was an opening pointing away from it, see Figure 2 for visual examples); and a condition in which the screen froze, but no lava was added/removed. Behavioral data showed that participants were much slower to respond in the last condition with no change, which may be because they were unsure if they had missed a subtle change in the environment (the lava appeared or disappeared instantaneously when they reached the change point). Thus, we focused our analysis on the other conditions. For Detours and Shortcuts, there were also two levels of change to the (optimal) new path, either 4 or 8 grid steps extra/less, respectively. See Figure 2 for example schematics of these changes. Finally, during the test phase there were also control ‘Follow’ trials which started with an arrow that indicated the direction to travel. In this case, participants were required to follow the twists and turns of the arrow until a new target object appeared, and from then onward the trial was like the ‘Navigation’ events as described above. The comparison of ‘Navigation’ vs ‘Follow’ movements allowed us to relate our results to those of previous experiments (Howard et al., 2014; Javadi et al., 2017; Patai et al., 2017).

### Spatial Parameters

Spatial parameters were calculated in the same manner as in Howard et al (2014) and Javadi et al (2017). In brief, Path Distance (PD), Euclidian Distance (ED), Egocentric Goal Direction (EGD) and the number of optimal upcoming Turns were calculated at each change point. All parameters were highly correlated (p<0.001, see Table 1A), except for PD/ED and EGD. Based on our previous work (Howard et al. 2014, Patai et al., 2017), our main analysis involved using PD as an independent parametric regressor (Table 1B). We also considered a control model that included both PD and EGD, as these measures were not correlated. The other parameters were not explored independently. Spatial parameter values were rescaled between 0 and 1, where 1 is the maximum value, e.g. the greatest distance.

**Table 1A:** Correlation between Spatial Parameters at Change Point / Start (Object onset). PD: new path distance after the change; PD%: relative change in path distance (compared to pre-change path distance); EGD: egocentric goal direction; ED: Euclidian distance, Turns: number of upcoming turns. Note PD% does not exist at the start of trial, i.e when the target object is presented, as this measure assumes a change from the original path, which is only available at Change Points. Shown are r values, with significance indicated by: **p<0.001; * p<0.05

|  | PD | PD% | ED | EGD | Turns |
| --- | --- | --- | --- | --- | --- |
| PD |  | 0.43** | 0.64**/0.07** | -0.03/-0.05* | 0.58**/0.56** |
| PD% |  |  | -0.1** | 0.09** | 0.49** |
| ED |  |  |  | -0.02/-0.07** | 0.11**/-0.11** |
| EGD |  |  |  |  | -0.22**/0.04 |

**Table 1B:** Details of GLM parameters for the fMRI models

| MODEL | CONDITIONS | PARAMETRIC REGRESSORS | DURATION |
| --- | --- | --- | --- |
| Terrain change type | Detours (+8/-8), Shortcuts (-8/-4), False Shortcuts (Away/Towards) | - | 4s |
| New Path at Change | All conditions together | PD &<br>PD+EGD (control) | 4s |
| Back-tracking (step-matched) | Back-tracking vs Non-Back-tracking | - | 0s |

**Figure 2:**
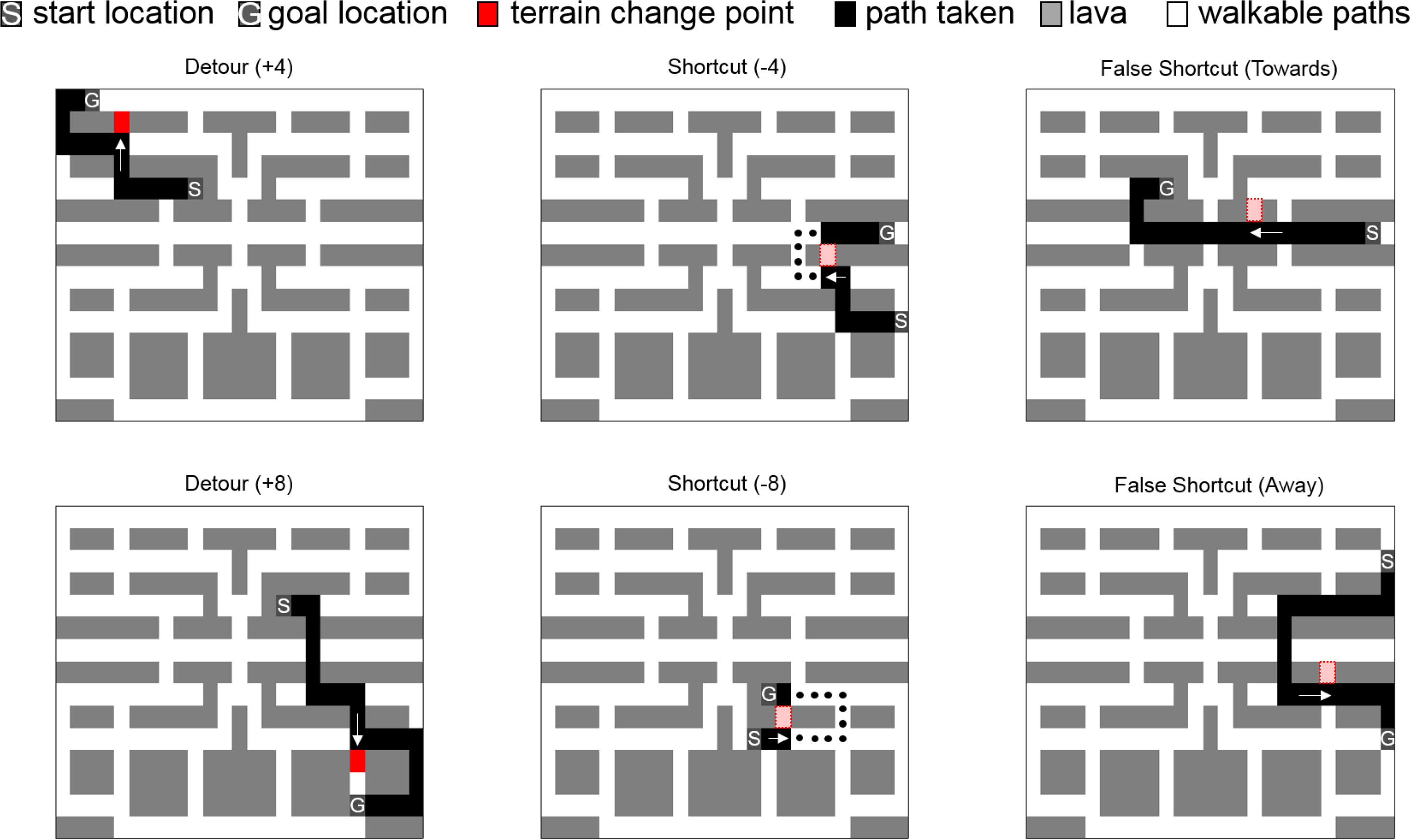
Examples of a changes that occurred during routes to the goal. Participants start their path from the last object they found and go towards the current (new) goal location along the shortest path available. For Detours, at some point along the route, the participant’s path is blocked and they are forced to take a detour around the lava to reach their goal. In the case of a shortcut, a grid point would be unblocked, thus revealing a novel, shorter route to the goal (originally optimal path shown in dots if no shortcut had been presented). In the case of False Shortcuts, taking this opening would be detrimental as it would lead to a longer path to the goal, despite the path seeming to head towards (or away) from it. The full grid was 25×15 squares, and is shown from above in these examples. +/− 4 or 8 refers to the amount added or subtracted in steps.

### fMRI Scanning & Preprocessing

Scanning was conducted at the Birkbeck-UCL Centre for Neuroimaging (BUCNI) using a 1.5 Tesla Siemens Avanto MRI scanner (Siemens Medical System, Erlangen, Germany) with a 12 channel head coil. Each experimental session lasted around 60 minutes and was separated in three parts (each of approximately 15-20 minutes). Approximately 980 functional scans were acquired per session (depending on routes taken), using a gradient-echo incremental EPI sequence (TR=3400ms, TE=50ms, TA=3.315s, flip angle= 90°). The slice thickness was 2mm with a gap of 1mm, TR=85ms, TE=50ms, slice tilt = 30°. The field of view was 192 mm, and the matrix size was 64 × 64. The scan was a whole brain acquisition, with 40 slices and a repetition time of 3.4 s. A T1-weighted high-resolution structural scan was acquired after the functional scans (TR=12ms, TE=5,6ms, 1×1×1mm resolution). Ear plugs were used for noise reduction, foam padding was used to secure the head in the scanner and minimize head movements. Stimuli were projected to the back screen, a mirror was attached to the head coil and adjusted for the subjects to see full screen. All fMRI preprocessing and analysis was performed using SPM12. To achieve T1 equilibrium, the first six dummy volumes were discarded. During preprocessing, we used the new Segment (with 6 tissue classes) to optimize normalization. Otherwise, we used all default settings, and we performed slice timing correction. No participants had any abrupt motion change over 4mm.

### Experimental Design and Statistical Analysis

Participants performed a total of 120 routes, with one change event occurring in each route (number of trials per condition was 17 on average, range: 11-25, depending on the different scenarios used for counterbalancing routes taken). Each route started from a previous goal and ended at the new goal object for that trial. We used repeated-measures ANOVAs to test for behavioural differences (accuracy, extra steps, back-tracking trials) between conditions. We also calculated d-prime and criterion (signal detection theory measures) to quantify the bias to take a False Shortcuts Towards instead of Away from a goal (both false alarms calculated relative to correct Shortcuts, which are hits). We recorded the response time to make the first choice after the 4 seconds elapsed, but due to the 4 second delay we do not interpret this as a traditional decision-making reaction time.

To analyse the fMRI data, we constructed multiple models, based on a priori predictions from previous work (Howard et al, 2014). Please see that Table 1B for a description of the models, events included and regression parameters. We used a standard preprocessing pipeline in SPM. A priori regions of interest were small volume corrected using anatomical masks (WFU Pick atlas [Maldjian, Laurienti, Kraft, & Burdette, 2003; Tzourio-Mazoyer et al., 2002]) and a functional mask for the dorsomedial PFC (Kaplan et al., 2017) was employed in one follow-up exploratory analysis. For completeness we also report all results at an uncorrected threshold of p < 0.001, minimum 5 contiguous voxels (Howard et al., 2014). This is provided to allow comparison to past similar datasets rather that to draw specific inferences about predicted responses. Note that we used all trials for an event type, irrespective of whether or not the participant was correct for not. We also report data for False Shortcuts comparing correct and incorrect choices in the Extended Data (Figure 3-1).

A ‘Back-tracking’ event was defined by when participants pressed the backwards button and returned to a step along the route they had just come down. Non-back-tracking events were selected from the participants’ other paths and were matched to these in the relative number of steps taken before the back-tracking happened (for example: halfway through the route). We did not use the absolute number of steps, because trials that contained a back-tracking event were often much longer, and thus the step number at which a back-tracking event occurred would in many cases already been located at (or past) a goal location in a non-back-tracking trial.

## Results

### Behaviour

Our primary measure of navigation was the accuracy of the whole route, in other words whether participants took the optimal path to the target. We conducted a repeated measures ANOVA to test for effects of terrain change type on participants’ accuracy in finding the correct path. We found that there was a significant effect of terrain change type (F(1,120)=17.7,p<0.001), such that Detours (+8) and False Shortcuts Towards the goal resulted in less optimal path taking (both t(1,20)<-3.6,p<0.002) compared to all other conditions (see Table 2, and Extended Data Table 1 for comprehensive t-tests).

To follow up the errors in which participants did not take the optimal path, we looked at the number of extra steps taken on a route. We found a significant effect of terrain change type (F(1,120)=8.3,p<0.001), with overall more steps off-route in the Detours (+8) and False Shortcuts Towards conditions. When quantifying the number of extra steps as a proportion of the total (new) number of optimal steps from the terrain change point onward for a given route, in fact Shortcuts resulted in the largest proportion off-route (see Table 2). Some of these extra steps were due to participants turning around, i.e., “Back-tracking”; these were again more common in the Detours (+8) condition (F(1,120)=8.5,p<0.001, compared to all t(1,20)>2.1, p<0.051, see Extended Data Table 2–6 for details). Overall, only 19% of the extra steps were such Back-tracking events (and this ratio was not significantly different between conditions, F(1,65)=2.17, p=0.068). Moreover, we calculated the ratio of correct, compared to incorrect, Back-tracking trials (“correct” is defined as a trials in which back-tracking would actually bring the participant closer to the goal), and found that overall 85% (±2% s.e.m, range: 50-100%) of Back-tracking events were correct or optimal, and occurred equally frequently across all conditions (F(1,5)=2.2,p=0.066). Thus, in the majority of Back-tracking events participants became aware that they were heading away from the goal and spontaneously decided to turn around.

**Table 2:** Behavioural Results Summary [mean (±s.e.m)]

|  | **Detour**<br>**(+8)** | **Detour**<br>**(+4)** | **Shortcut**<br>**(-8)** | **Shortcut**<br>**(-4)** | **False**<br>**Shortcuts**<br>**Towards** | **False**<br>**Shortcuts**<br>**Away** |
| --- | --- | --- | --- | --- | --- | --- |
| **Accuracy [%]** | 64.1(±3.9) | 80(±2.3) | 84.6(±2.6) | 84.6(±2.8) | 65.8(±3.1) | 81.3(±1.9) |
| **Extra Steps** | 1.9(±0.26) | 0.99(±0.14) | 1.5(±0.37) | 1.2(±0.3) | 2(±0.31) | 0.97(±0.17) |
| **Extra Steps**<br>**[proportion]** | 1.2(±0.11) | 1.4(±0.6) | 2.1(±1.07) | 1.7(±0.37) | 1.3(±0.18) | 1.4(±0.32) |
| **Back-tracking [#]** | 7.1(±1.3) | 3.7(±0.7) | 2.2(±0.6) | 2.6 (±0.6) | 4.3(±0.6) | 3.5(±0.4) |
| **Back-tracking**<br>**[% of extra]** | 22(±5.5) | 19(±2.4) | 19(±4.4) | 22(±6) | 14(±2.6) | 29(±3.8) |
| **Correct**<br>**Backtracking [%]** | 85(±3.1) | 83(±3.7) | 78(±5.5) | 70(±7.8) | 80(±4.5) | 71(±7.7) |

### fMRI Results

fMRI analyses revealed that bilateral hippocampus, posterior cingulate and retrosplenial cortex, as well as frontal areas were more active when participants were actively navigating than when they merely followed an arrow on the screen (Extended Data Table 7). Both the left and the right hippocampus were significantly more active in the navigate than the follow condition (small-volume correction, FWE p<0.05), in line with previous findings (Howard et al., 2014; Patai et al., 2017, for an overview see Spiers & Gilbert 2015).

#### Caudate, but not hippocampal, activity responds to changes in the path to the goal at Detours

We predicted that hippocampal activity would track the parametric change in the path distance to the goal when the structure of the environment changed. We found no evidence to support this prediction, both with specific ROIs and at a low uncorrected threshold (p < 0.005). This was also true when large Detours (+8) were directly compared with small Detours (+4). By contrast, we found that activity in the caudate nucleus bilaterally tracked the change in the path distance across all types of events (see Table 3), complementing past evidence that this region tracked the magnitude of change in the path at Detours (Howard et al., 2014).

#### Superior and right lateral prefrontal cortex respond to preferentially to Detours

Next we investigated the brain areas that have been found in previous studies when comparing Detours to non-Detours, including the superior frontal gyrus (SFG), the right lateral prefrontal (rlPFC) and frontopolar cortex (Spiers & Gilbert, 2015), using a combined mask of these areas. A linear contrast of terrain change type: Detours (+8) > Detour (+4) > Shortcuts (−4) > Shortcuts (−8), revealed a significant effect, with specifically the SFG and rlPFC activity scaling with the deviation from the optimal path prior to the change (Figure 3A and Table 3). False Shortcuts Towards the goal also significantly activated the rlPFC compared to Shortcuts (Figure 3B, and Table 3). By contrast False Shortcuts Away from the goal did not drive activity in rlPFC. Frontopolar cortex was not found to be modulated by Detours, Shortcuts, or False Shortcuts.

#### Prefrontal responses evoked during spontaneous internally generated Back-tracking events

We were interested in the brain areas that were activated by spontaneous, internally generated route changes, i.e. Back-tracking events. To investigate their neural correlates, we compared moments in routes where a back-tracking event occurred (defined as a return to a previous grid point along a single journey towards a target), to equivalent events where no back-tracking happened (equalized according to relative steps along a journey). When comparing Back-tracking to non-Back-tracking events, we again found significant SFG and rlPFC activations (Figure 4A, and Table 3). These results suggest that the prefrontal regions previously found to be involved in detour processing are also active when route changes occur due to spontaneous, internally generated events. We did not find evidence that the frontopolar cortex was responsive to Back-tracking.

Outside our ROIs drawn from Spiers and Gilbert (2015), we found that during Back-tracking, activity in a region of medial prefrontal cortex survived whole-brain FWE correction (Figure 4). This activation overlapped with the dorsal anterior cingulate area (dACC) reported by a recent study when difficult navigational decisions were required (Kaplan et al., 2017). We used a functional ROI (includes the dACC and supplementary motor area (SMA)) from Kaplan et al. (2017) to investigate whether this area was also active during terrain change points, and found that for both the linear contrast above (Detours (+8) > Detour (+4) > Shortcuts (−4) > Shortcuts (−8)) and False Shortcuts Towards > Shortcuts contrast this area was significantly active (Table 3). However when plotting the peak activation, we found that Backtracking events resulted in a significantly larger activation of the dACC than the other contrasts (both t>5.4,p<0.001; Figure 4, bottom right), pointing to a potentially unique role during the processing of internally generated route changes, which presumably occurs when participants become aware their current route plan is not effective.

#### Back-tracking events disengage from the putative Default-Mode Network

Because Back-tracking involved a visual change in view (180°) compared to Non-back-tracking events we explored whether similar regions would be active when participants turned left or right compared to when they did not turn. We found a significant response in the functional dACC mask when comparing Turns to Non-turns, however the peak was located in the SMA. When Back-tracking was compared to Turns the dACC was significantly more active in Back-tracking events, in which an equivalent amount of visual change is contrasted (90° difference in change of viewpoint). Conversely, Turns compared to Back-tracking resulted in significant responses in the hippocampus, anterior medial PFC and posterior cingulate cortex. These regions are overlapping with those implicated in the default-mode network (DMN) (Andrews-Hanna, Reidler, Sepulcre, Poulin, & Buckner, 2010). To explore whether the results match the default-mode network more explicitly we created an aggregate ROI including the medial PFC, the precuneus/posterior cingulate cortex, bilateral parahippocampal cortex and angular gyrus. We found that Back-tracking significantly suppressed this putative DMN compared with Turns, and this was also the case when comparing Back-tracking to Non-back-tracking and Detour events, highlighting disengagement from this network when participants spontaneously instigated a route change (Figure 5).

#### Control Analysis for Route Visibility

As our maze environment was open-plan, and participants could see ahead (see Figure 1, ‘Test’), it should be noted that the task could have been solved using a purely visual search of the available paths to the remembered goal location during the change period (as opposed to relying on the map of the layout of the environment from memory). In this case we would expect that at change points where visual analysis of the scene is restricted due to the heterogeneity of the maze layout from the point where they are stopped, to be lower than when they can ‘look’ ahead to check the available paths to the goal. This is particularly relevant for False Shortcuts where ideally participants would decide whether to take the opening based on pre-existing knowledge of the maze layout and the connecting paths to decide if it is a real shortcut or not. We compared False Shortcuts Towards the goal in two cases: when the upcoming path was clearly visible from the change point and when it was not, and found a significant effect in accuracy (visible: 76%(±4), not clearly visible: 53%(±5)), however we also note that in the latter instances, it was the case that taking the False Shortcut only resulted in a minor addition of steps to the path (maximum 3 steps), whereas when then alternative path was more visible, taking those false entry points would have resulted in a significant lengthening to the path to the goal (10+ steps). Due to trial numbers, we cannot directly compare visible vs non-visible by accuracy. Based on these results we cannot clearly confirm whether or not participants were solely using a cognitive map to solve the problem at change points, or if they were also engaging in visual search behaviour. Importantly, peak activity in the right lateral PFC activity seen in the contrast of False Shortcut Towards vs Shortcuts (Figure 3B), was not significantly different between ‘visible’ and ‘not clearly visible’ False Shortcuts Towards the goal (p>0.1), underscoring the notion that in order to correctly reject a false opening, participants did engage prefrontal areas.

**Table 3:**
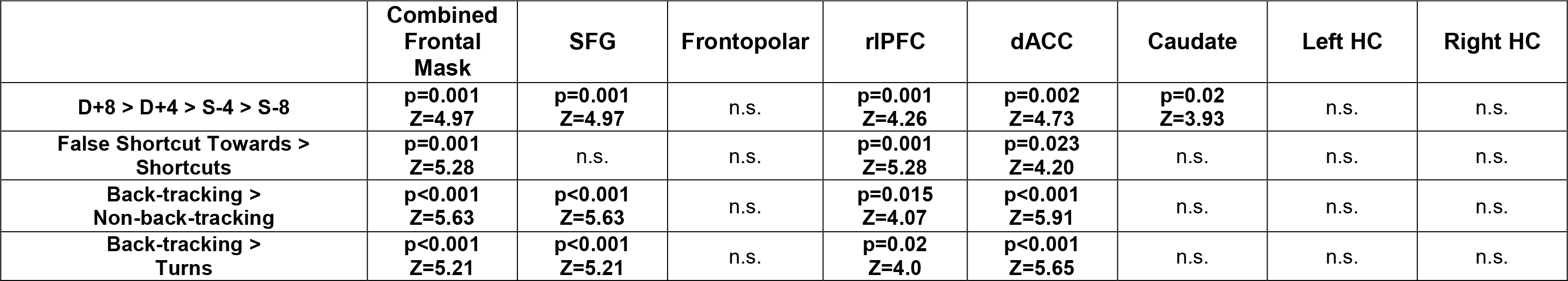
Results of small-volume correction in ROIs during Terrain Changes and Back-tracking. (All results reported are significant after FDR correction)

|  | **Combined**<br>**Frontal**<br>**Mask** | **SFG** | **Frontopolar** | **rIPFC** | **dACC** | **Caudate** | **Left HC** | **Right HC** |
| --- | --- | --- | --- | --- | --- | --- | --- | --- |
| **D+8 > D+4 > S-4 > S-8** | **p=0.001**<br>**Z=4.97** | **p=0.001**<br>**Z=4.97** | n.s. | **p=0.001**<br>**Z=4.26** | **p=0.002**<br>**Z=4.73** | **p=0.02**<br>**Z=3.93** | n.s. | n.s. |
| **False Shortcut Towards >**<br>**Shortcuts** | **p=0.001**<br>**Z=5.28** | n.s. | n.s. | **p=0.001**<br>**Z=5.28** | **p=0.023**<br>**Z=4.20** | n.s. | n.s. | n.s. |
| **Back-tracking >**<br>**Non-back-tracking** | **p<0.001**<br>**Z=5.63** | **p<0.001**<br>**Z=5.63** | n.s. | **p=0.015**<br>**Z=4.07** | **p<0.001**<br>**Z=5.91** | n.s. | n.s. | n.s. |
| **Back-tracking >**<br>**Turns** | **p<0.001**<br>**Z=5.21** | **p<0.001**<br>**Z=5.21** | n.s. | **p=0.02**<br>**Z=4.0** | **p<0.001**<br>**Z=5.65** | n.s. | n.s. | n.s. |

**Figure 3:**
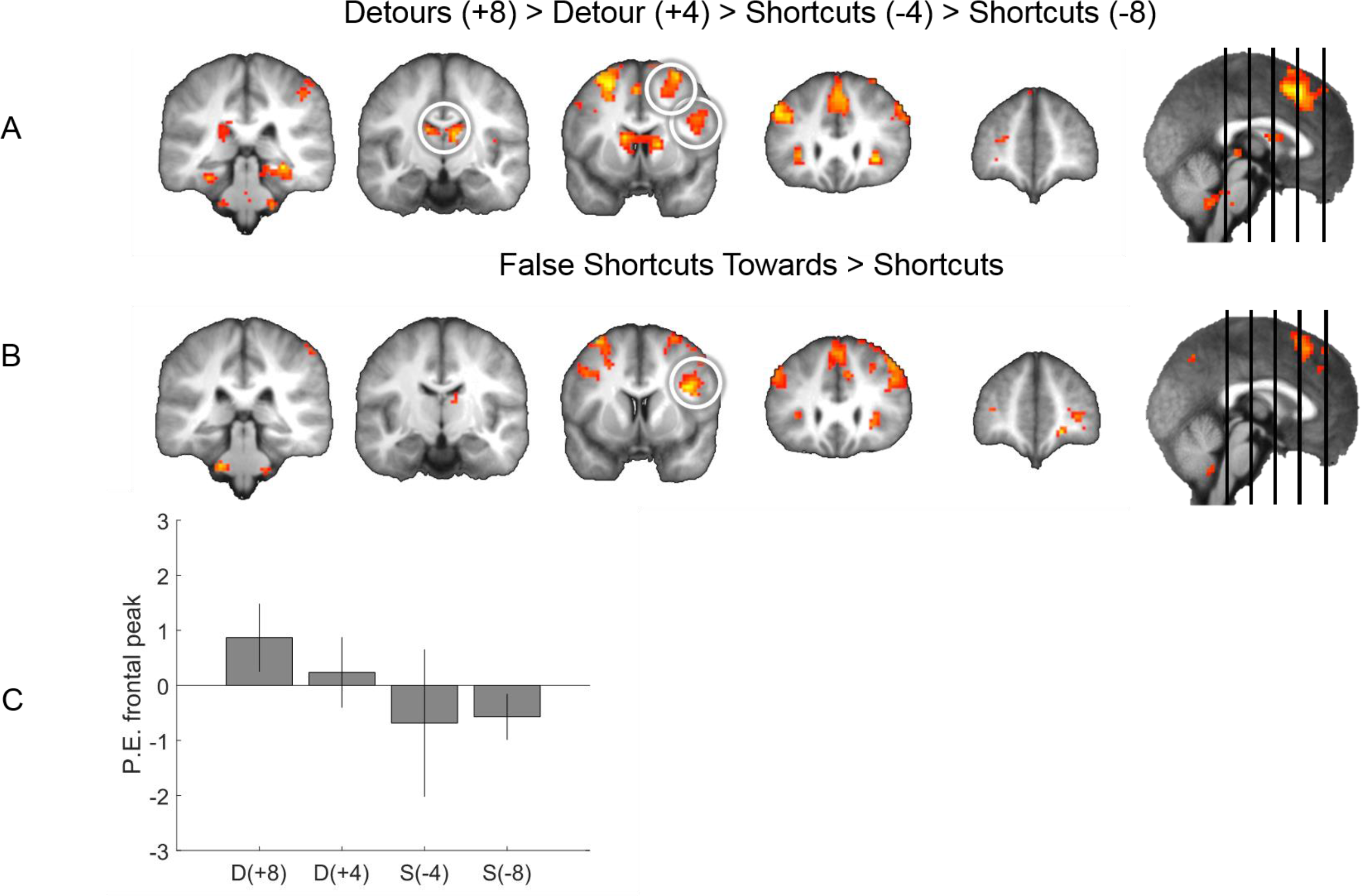
Prefrontal Areas involved during processing of Terrain Changes. A) Superior frontal gyrus and right lateral PFC were engaged in the linear contrast of Detours and Shortcuts [Detours (+8) > Detour (+4) > Shortcuts (−4) > Shortcuts (−8)] and B) right lateral PFC when comparing False Shortcuts Towards the goal to Shortcuts. Purely for display purposes, figures are thresholded at p=0.005 uncorrected, minimum 5 contiguous voxels. Significance was assessed using voxel-wise corrected ROIs C) Parameter estimates from the peak frontal voxel in the linear contrast from A, to illustrate the effect. Figure 3-1 contains the contrast of correct vs incorrect choices at False Shortcuts Towards the goal.

**Figure 4:**
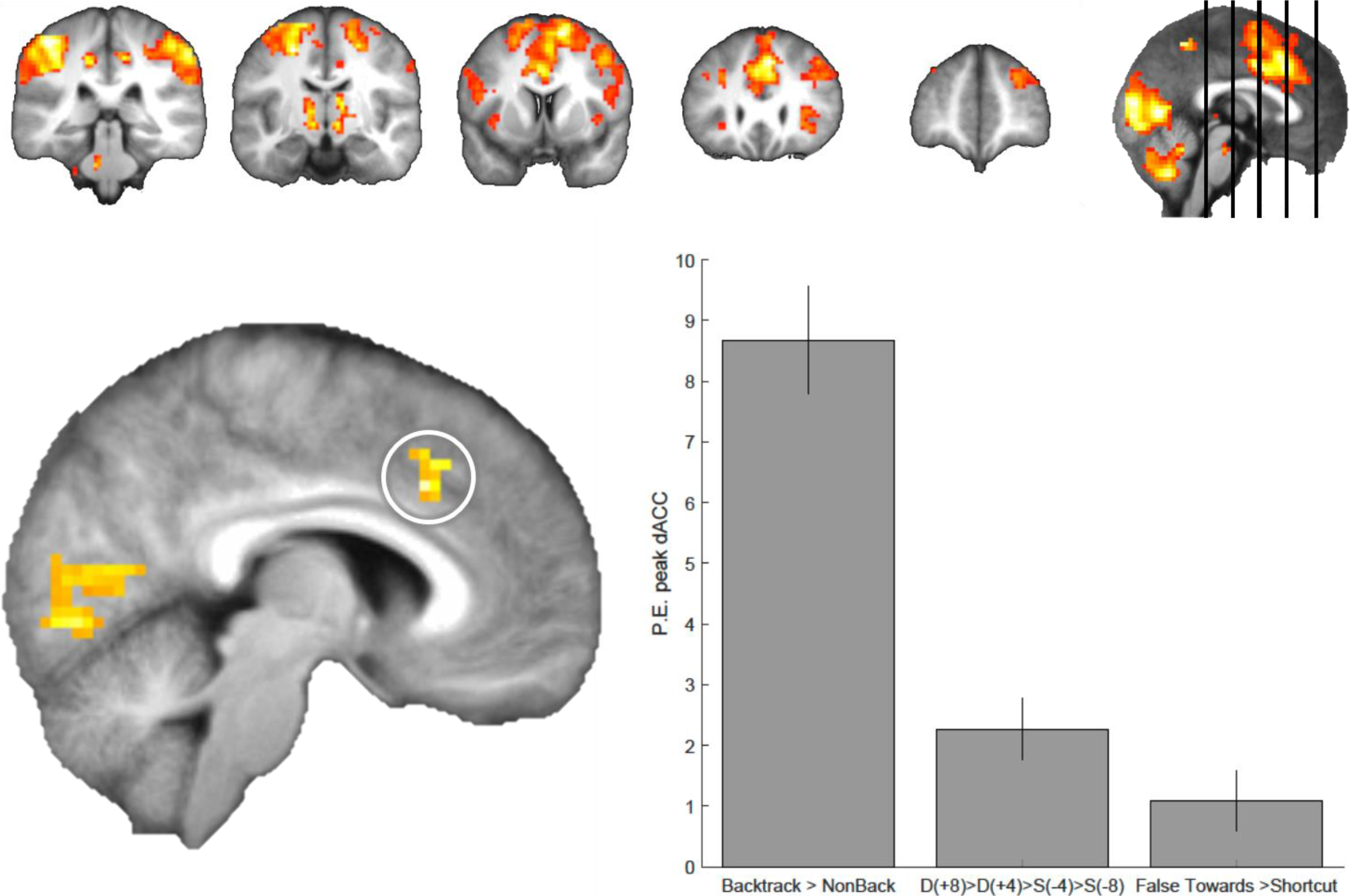
Back-tracking activates a range of frontal areas, as well as the dorsal anterior cingulate cortex. Top: Similarly to Detours, the superior frontal gyrus and right lateral PFC are activated during Back-tracking compared to Non-back-tracking events. Figures are thresholded at p=0.005 uncorrected. Bottom: Whole-brain results (FWE p<0.05) revealed a significant activation in the dorsal anterior cingulate cortex. Extracting parameter estimates from the peak voxel in dACC [MNI: x:6 y:20 z:35, note Kaplan et al. (2017) = x:6 y:23 z:37] show that Back-tracking and Detours activate this region significantly, however Back-tracking activates it significantly more than Detours and False Shortcuts Towards the goal.

**Figure 5:**
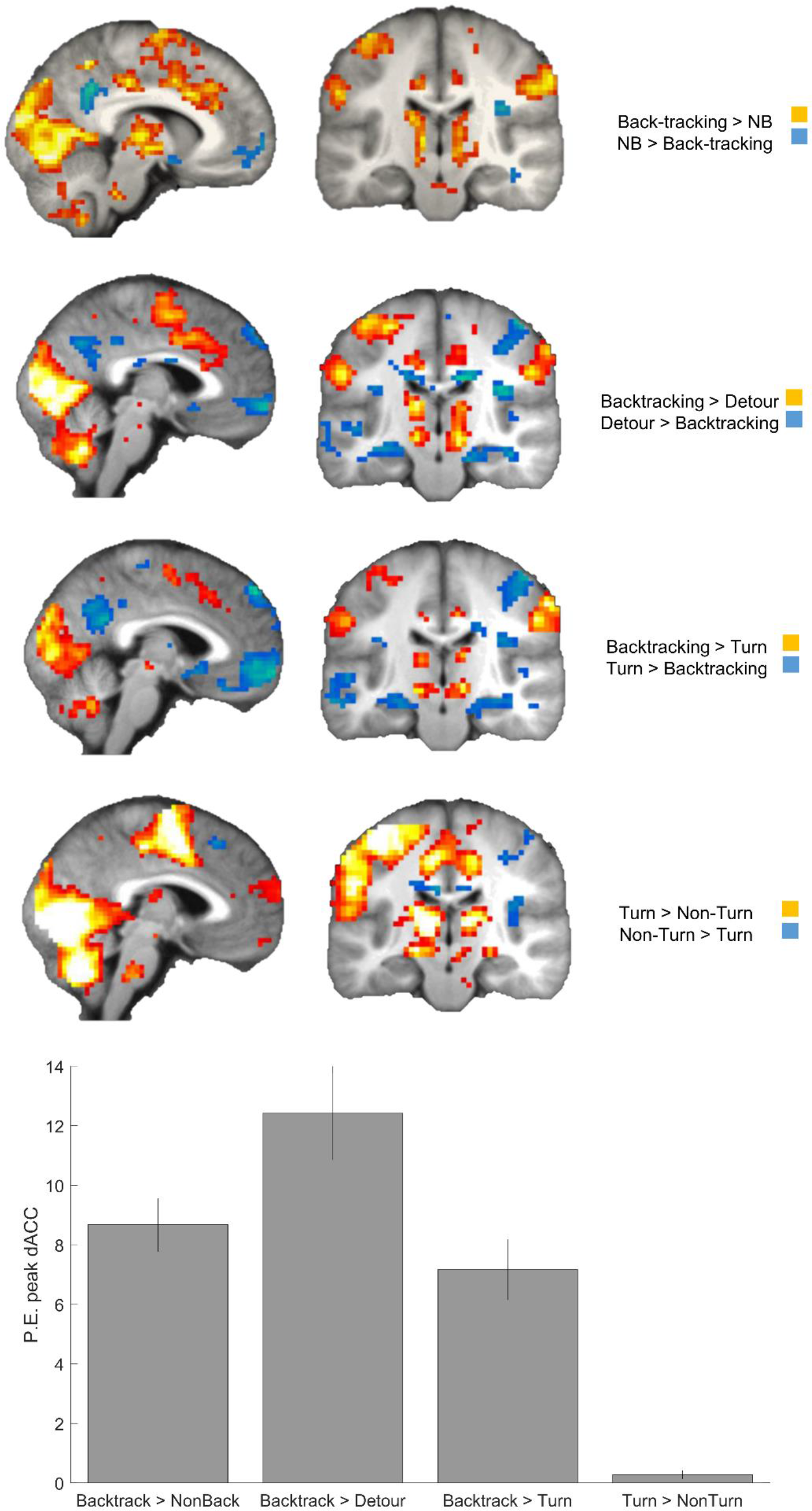
Back-tracking events disengage regions in the putative Default-mode network when compared to other events. Results are shown at p=0.005 uncorrected, minimum 5 contiguous voxels. Significant results in the combined DMN mask: Backtracking compared to Non-backtracking (p=0.038, Z=4.21), Turns (p=0.006, Z=4.62) and Detours (p=0.008, Z=4.55).

## Discussion

A core tenet of the cognitive map theory is that internal representations support flexible navigation, enabling an animal to make use of shortcuts and take efficient detours (O’Keefe & Nadel, 1978; Tolman, 1948). Despite the wide acclaim for this idea, little research, especially in humans, has been directed at the neural mechanisms of such adaptive behaviour (Spiers and Gilbert, 2015; Epstein et al., 2017). Using fMRI and a VR task involving navigation through a landscape that changed layout sporadically, we examined the neural responses to forced Detours, novel Shortcuts, False Shortcuts and Back-tracking. We found: i) superior and lateral PFC and caudate activity responded to Detours, maximally when there was a large change in the path, i) rlPFC responded when false shortcuts to the goal needed to be avoided and iii) a sub-region of anterior cingulate increased its activity during spontaneous Back-tracking in the context of de-activation of the regions associated with the default-mode network.

### The role of prefrontal cortex, hippocampus and caudate in responding to detours and shortcuts

Based primarily on evidence from nine fMRI studies, Spiers & Gilbert (2015) provided preliminary predictions about how the PFC and the hippocampus might respond to forced detours and changes in the layout of an environment. Lateral PFC was suggested to provide a prediction error signal in response to changes in the path options (responding whenever an unpredicted change in the possible paths occurs). The superior and anterior PFC were speculated to support re-formulation of the route plan (responding at all events that require reconsidering the change in route plan). The hippocampus was postulated to simulate the future path the goal (responding the greater the increase in the path to the goal), drawing on rodent place cell studies (Ólafsdóttir, Barry, Saleem, Hassabis, & Spiers, 2015; Pfeiffer & Foster, 2013). Here we failed to find evidence that the hippocampus specifically encodes the change in the path distance to the future goal. One possibility is that the hippocampus simulates future possible scenes (Hassabis & Maguire, 2007), re-constructing the different locations that lie between the current location and the future goal (Javadi et al., 2017; Spiers & Barry, 2015). In the case of the current study the environment was sparse with few features to distinguish different parts of the island, which might explain why we did not observe a correlation between the hippocampus and the change in path to the goal. Notably, previous studies reporting hippocampal activity correlated with the future path to the goal used real-world stimuli with nameable landmarks located along the paths (Howard et al., 2014; Javadi et al., 2017; Patai et al., 2017).

By contrast to the hippocampus, we found that activity in lateral and superior PFC, as well as the caudate, responded maximally when there was a large change in the path to the goal. The caudate response is consistent with a prior result from Howard et al. (2014) which found that the larger the detour the more activity was elicited in the caudate nucleus. Thus, speculatively the caudate activity may relate to a signal linked to updating the transition structure in the environment at that particular location where the change occurs. An alternative is that the caudate plays an important role in updating its representations in relation to a model of the environment, consistent with this region coding a prediction error about future events (O’Doherty et al., 2004). Consistent with our caudate responses reflecting a model-based updating process, a previous fMRI study of navigation in a continually changing environment found that caudate activity correlated with parameters of a model-based representation of the environment (Simon & Daw, 2011).

The PFC responses we observed are in agreement with the predicted roles of the superior PFC supporting resolving path conflict and the rlPFC processing a prediction error signal between the predicted state of the world and the encountered layout (Spiers & Gilbert, 2015). Two types of prediction error could be processed in the current paradigm. One is the signed prediction error signal linked to the difference in the path before and after the change in the layout (+ve for detours, -ve for shortcuts). The other is an unsigned prediction error where the amount of change is coded rather than the direction of change (+ve for both detours and shortcuts). Our results show a wide network of regions including our PFC and caudate ROIs were driven in a manner consistent with the signed prediction error (maximal for +8 Detours). Our results thus align more strongly with models in which the PFC and caudate code the increase in path, and rather than being driven in a clear linear manner by the signed prediction error, the data suggest these regions might be driven in a threshold manner by large detours over the other conditions. Future research carefully varying along a broader range the amount of path change at detours will be required to explore these possibilities.

It is possible the PFC responses to Detours are driven by the presence of the physical barrier appearing to block the route. This is certainly a possibility in several past studies (e.g. Iaria et al., 2008; Maguire et al., 1998), though not all (see Howard et al., 2014) However, because rlPFC was more active for false shortcuts compared to shortcuts and these two events are visually similar (one unit of lava is removed to create a new path), it seems rlPFC is driven by planning demands rather than the visual processing of a barrier. The response is consistent with it playing a role in behavioural control: suppressing the pre-potent response to move towards the goal drawing on the observation that there is a now a barrier or that there is a new opening that is not helpful (Spiers & Gilbert, 2015).

### Spontaneous Internally Driven Changes in Route - Backtracking

Across both externally driven (Detours, False Shortcuts) and internally driven (Backtracking) route re-evaluation events we found increased activation in the dorso-medial PFC, an area that has been implicated in various contexts including: hierarchical planning (Balaguer, Spiers, Hassabis, & Summerfield, 2016), high planning demand decisions (Kaplan et al., 2017), and model updating, irrespective of difficulty or simple error-related signalling (Kolling et al., 2016; O’Reilly et al., 2013). However, we found activation in the dorsal anterior cingulate cortex (dACC) was much more pronounced during Back-tracking, possibly reflecting stronger engagement of planning and error signals when participants spontaneously realized that they were on a sub-optimal path. This is further supported by the result that nearly 80% of Back-tracking events were usefully corrective, resulting in a shorter path to the goal. One previous study also found medial PFC to be correlated with a model-based estimate of behaviour relating to putative events of back-tracking in a simple maze (Yoshida & Ishii, 2006), and our peak activation for the dACC is highly proximal to the one previously reported in that study. This highlights the role of the medial PFC/dorsal anterior cingulate in situations in which re-evaluation and updating from internal monitoring are required, as is the case during introspective awareness (Critchley, Wiens, Rotshtein, Öhman, & Dolan, 2004), and reorienting and error detection (Wang, Ulbert, Schomer, Marinkovic, & Halgren, 2005)(Carter et al., 1998). It remains to be tested whether the underlying error signal comes from the medial PFC/dACC itself or alternatively this area is involved in orchestrating the reorienting of attentional and memory systems based on incoming uncertainty signals.

Additionally, when comparing Back-tracking to Non-back-tracking, Detours or Turn events, we found a disengagement from the putative default-mode network (including the medial PFC, hippocampi, posterior cingulate cortex), with an increase in right lateral-frontal activity, consistent the notion that internally generated route changes resulted in a global resetting of attention from internal to external sources (Sridharan, Levitin, & Menon, 2008). The default-mode network has been implicated during transitions between tasks/states and at task restarts (Crittenden, Mitchell, & Duncan, 2015; Fox, Snyder, Barch, Gusnard, & Raichle, 2005; Smith, Mitchell, & Duncan, 2018), when participants were ‘in the zone’ during a self-generated behaviour (Kucyi, Hove, Esterman, Hutchison, & Valera, 2017), and before an error is committed (Eichele et al., 2008). Given that Backtracking involves a spontaneous realization of an incorrect path and an update in route plans, reductions in default-mode network activity and activation of the frontal areas including the ACC imply that this behaviour is similar to scenarios in which novel stimuli or task rules activate the saliency network (Seeley et al., 2007). It will be useful in future research with navigation and such non-navigation paradigms to dissociate making novel responses from the novel responses which require an adjustment in the plan for future action.

## Conclusion

Overall, our results help evaluate putative models of how the brain responds during back-tracking, shortcuts and detours. Future studies separating conflict between route options from prediction errors, search processes and updating representations will be needed to better understand how the brain adapts to changing environments. More importantly, exploring spontaneous human behaviour during naturalistic tasks such as navigation, including error monitoring and updating, could lead to more applications for cross-species comparisons of general problem-solving behaviour.

## Extended Data

**Figure 3-1:**
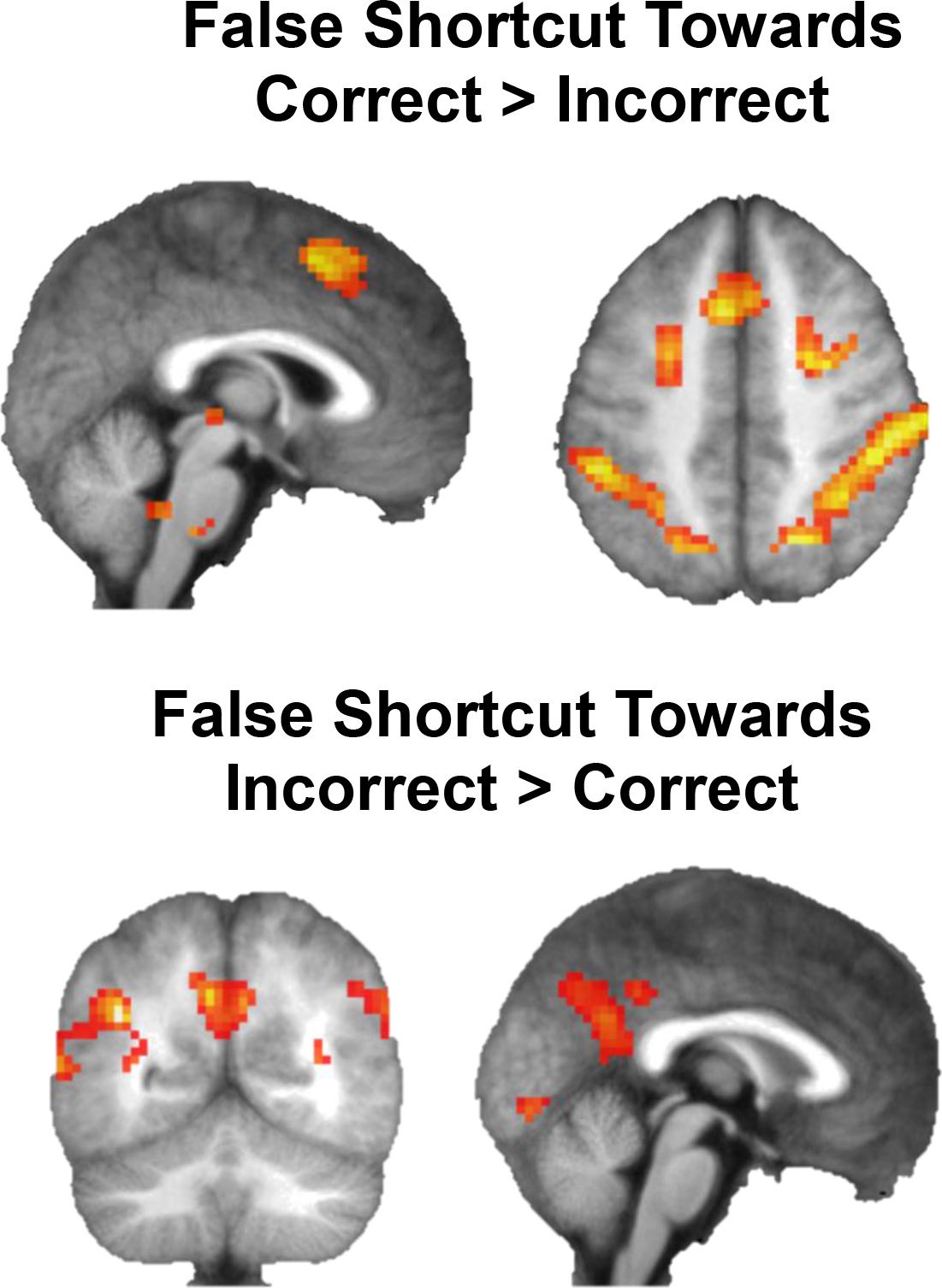
Contrast of Correctly Rejected vs Incorrectly Taken False Shortcuts. When participants correctly rejected the False Shortcut, there was more fronto-parietal activity, versus more visual and posterior cingulate activity when they chose incorrectly. Results are shown at p=0.005 uncorrected, minimum 5 contiguous voxels.

**Table S1:** Paired Samples T-Test comparing all terrain change types: Accuracy

|  |  | t | df | p |
| --- | --- | --- | --- | --- |
| **Detour (+8)** | - Detour (+4) | **-4.210** | **20** | **< .001** |
| **Detour (+8)** | - Shortcut (-8) | **-5.659** | **20** | **< .001** |
| **Detour (+8)** | - Shortcut (-4) | **-5.858** | **20** | **< .001** |
| **Detour (+8)** | - False Shortcut away | **-4.507** | **20** | **< .001** |
| Detour (+8) | - False Shortcut towards | -0.442 | 20 | 0.663 |
| Detour (+4) | - Shortcut (-8) | -1.468 | 20 | 0.158 |
| Detour (+4) | - Shortcut (-4) | -1.375 | 20 | 0.184 |
| Detour (+4) | - False Shortcut away | -0.487 | 20 | 0.632 |
| Detour (+4) | - **False Shortcut towards** | **3.655** | **20** | **0.002** |
| Shortcut (-8) | - Shortcut (-4) | -0.012 | 20 | 0.990 |
| Shortcut (-8) | - False Shortcut away | 1.047 | 20 | 0.307 |
| Shortcut (-8) | - **False Shortcut towards** | **5.059** | **20** | **< .001** |
| Shortcut (-4) | - False Shortcut away | 1.068 | 20 | 0.298 |
| Shortcut (-4) | - **False Shortcut towards** | **6.197** | **20** | **< .001** |
| False Shortcut away | - **False Shortcut towards** | **5.515** | **20** | **< .001** |

Note. Student's T-Test. Bold indicates significance after Bonferroni correction.

**Table S2:**
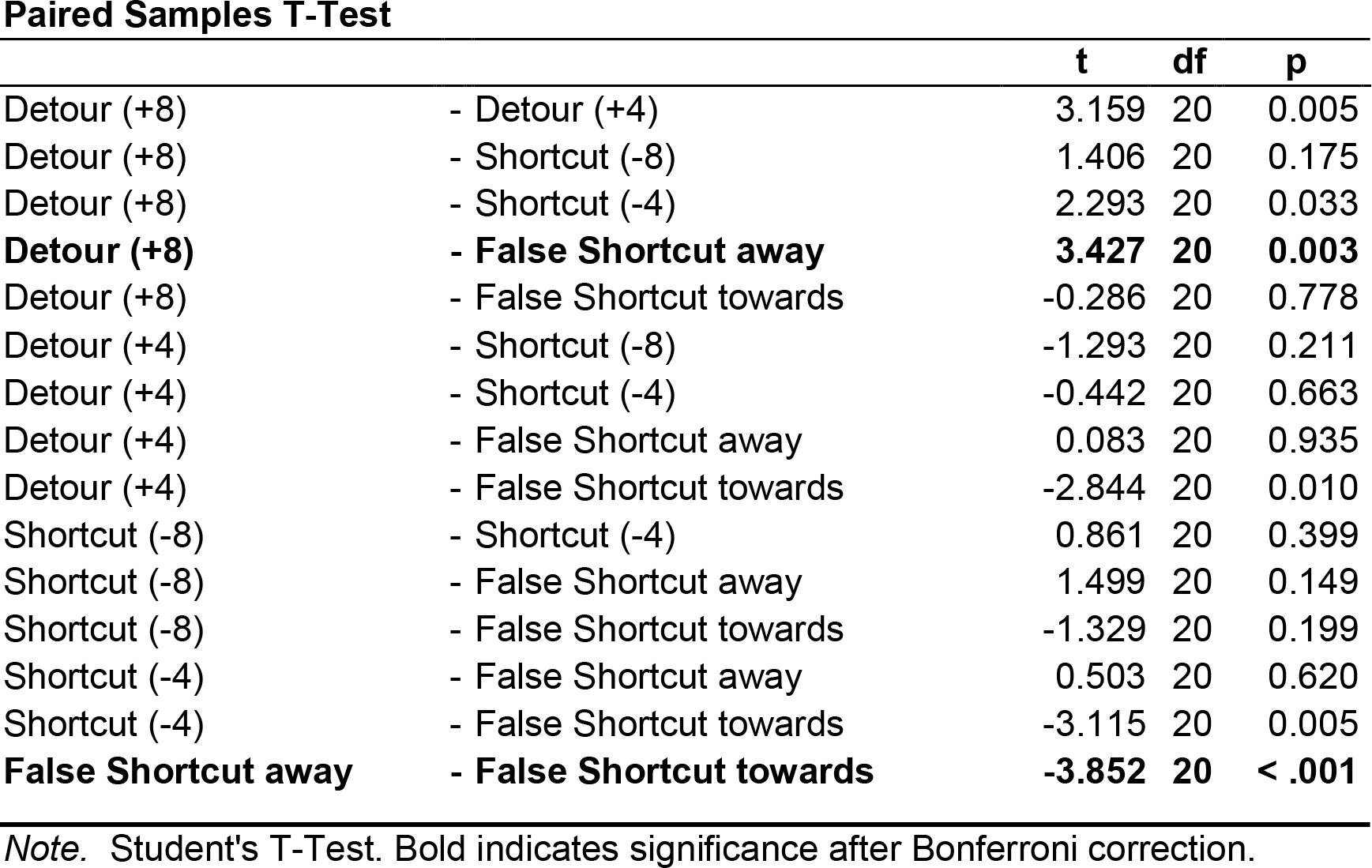
Paired Samples T-Test comparing all terrain change types: Extra steps

**Table S3:**
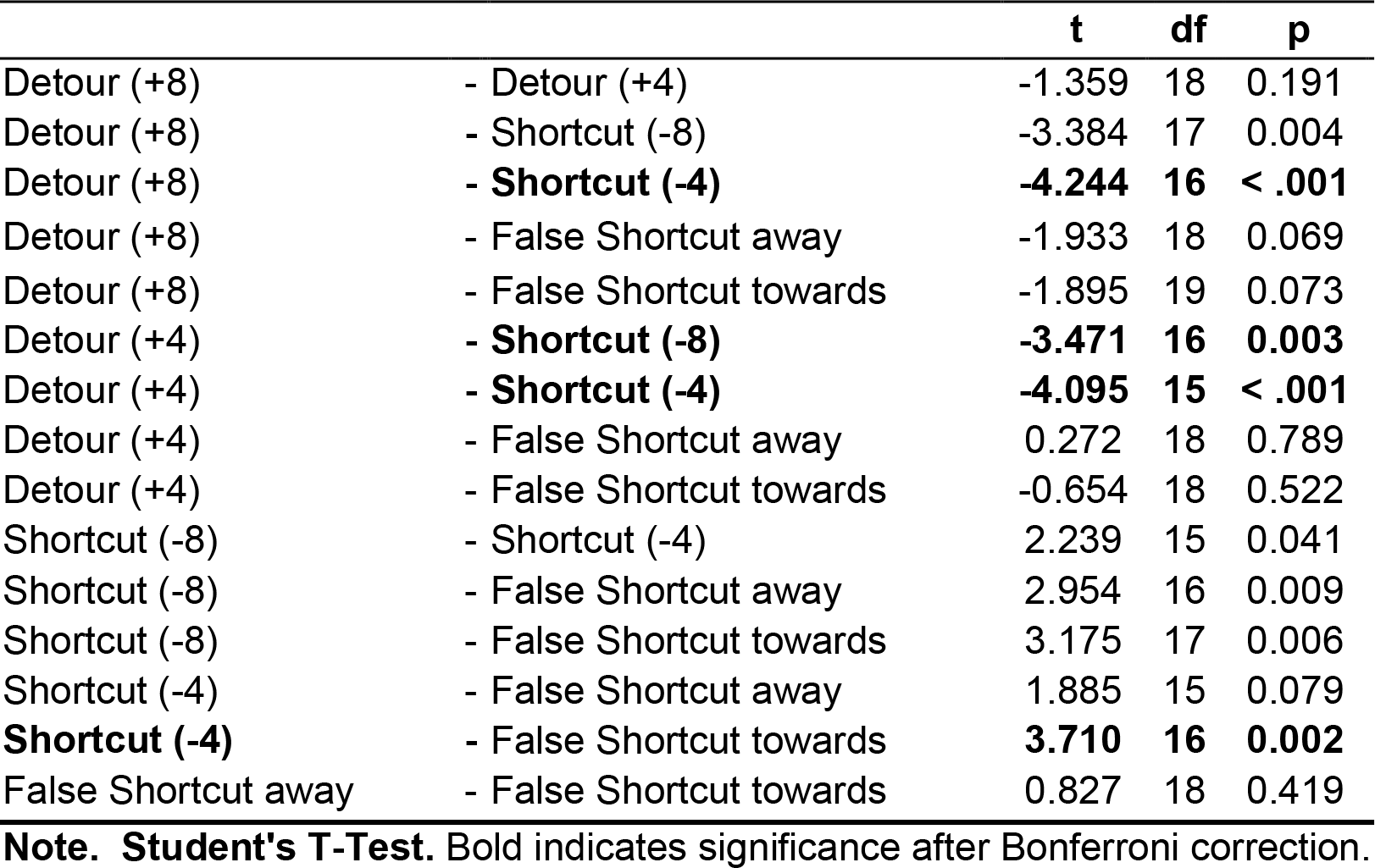
Paired Samples T-Test comparing all terrain change types: Extra steps proportions

**Table S4:** Paired Samples T-Test comparing all terrain change types: Back-tracking occurrences

|  |  | t | df | p |
| --- | --- | --- | --- | --- |
| **Detour (+8)** | - Detour (+4) | 2.548 | 20 | 0.019 |
| **Detour (+8)** | - Shortcut (-8) | 4.458 | 20 | < .001 |
| **Detour (+8)** | - Shortcut (-4) | 3.696 | 20 | 0.001 |
| **Detour (+8)** | - False Shortcut away | 2.759 | 20 | 0.012 |
| **Detour (+8)** | - False Shortcut towards | 2.073 | 20 | 0.051 |
| Detour (+4) | - Shortcut (-8) | 1.786 | 20 | 0.089 |
| Detour (+4) | - Shortcut (-4) | 1.884 | 20 | 0.074 |
| Detour (+4) | - False Shortcut away | 0.386 | 20 | 0.704 |
| Detour (+4) | - False Shortcut towards | -0.669 | 20 | 0.511 |
| Shortcut (-8) | - Shortcut (-4) | -0.559 | 20 | 0.583 |
| Shortcut (-8) | - False Shortcut away | -1.989 | 20 | 0.061 |
| Shortcut (-8) | - **False Shortcut towards** | **-2.350** | 20 | 0.029 |
| Shortcut (-4) | - False Shortcut away | -1.579 | 20 | 0.130 |
| Shortcut (-4) | - False Shortcut towards | -2.024 | 20 | 0.056 |
| False Shortcut away | - False Shortcut towards | -1.116 | 20 | 0.278 |

**Note. Student's T-Test.** Bold indicates significance after Bonferroni correction.

**Table S5:** Paired Samples T-Test comparing all terrain change types: Back-tracking in relation to overall extra steps

|  |  | t | df | p |
| --- | --- | --- | --- | --- |
| Detour (+8) | - Detour (+4) | 0.824 | 18 | 0.421 |
| Detour (+8) | - Shortcut (-8) | 1.475 | 15 | 0.161 |
| Detour (+8) | - Shortcut (-4) | 0.155 | 16 | 0.879 |
| Detour (+8) | - False Shortcut away | -1.057 | 18 | 0.305 |
| Detour (+8) | - False Shortcut towards | 1.448 | 18 | 0.165 |
| Detour (+4) | - Shortcut (-8) | 0.066 | 16 | 0.948 |
| Detour (+4) | - Shortcut (-4) | -0.509 | 17 | 0.618 |
| Detour (+4) | - False Shortcut away | -2.398 | 18 | 0.028 |
| Detour (+4) | - False Shortcut towards | 1.319 | 19 | 0.203 |
| Shortcut (-8) | - Shortcut (-4) | -0.295 | 15 | 0.772 |
| Shortcut (-8) | - False Shortcut away | -1.565 | 15 | 0.138 |
| Shortcut (-8) | - False Shortcut towards | 1.125 | 16 | 0.277 |
| Shortcut (-4) | - False Shortcut away | -0.657 | 16 | 0.520 |
| Shortcut (-4) | - False Shortcut towards | 1.573 | 17 | 0.134 |
| **False Shortcut away** | **- False Shortcut towards** | **4.005** | **18** | **< .001** |

**Note.** Student's T-Test. Bold indicates significance after Bonferroni correction.

**Table S6:** Paired Samples T-Test comparing all terrain change types: Percentage of ‘Correct’ Back-tracking Events

|  |  | t | df | p |
| --- | --- | --- | --- | --- |
| Detour (+8) | - Detour (+4) | 1.503 | 21 | 0.148 |
| Detour (+8) | - Shortcut (-8) | 0.941 | 17 | 0.360 |
| **Detour (+8)** | - Shortcut (-4) | 2.343 | 21 | 0.029 |
| Detour (+8) | - False Shortcut away | 1.592 | 17 | 0.130 |
| Detour (+8) | - False Shortcut towards | 1.724 | 21 | 0.099 |
| Detour (+4) | - Shortcut (-8) | 0.362 | 17 | 0.722 |
| Detour (+4) | - Shortcut (-4) | 2.020 | 21 | 0.056 |
| Detour (+4) | - False Shortcut away | 1.259 | 17 | 0.225 |
| Detour (+4) | - False Shortcut towards | 1.073 | 21 | 0.296 |
| Shortcut (-8) | - Shortcut (-4) | 1.902 | 17 | 0.074 |
| Shortcut (-8) | - False Shortcut away | -1.394 | 17 | 0.181 |
| Shortcut (-8) | - False Shortcut towards | -0.638 | 17 | 0.532 |
| Shortcut (-4) | - False Shortcut away | -1.983 | 21 | 0.061 |
| Shortcut (-4) | - False Shortcut towards | -1.279 | 17 | 0.218 |

**Note.** Student's T-Test. Bold indicates significance after Bonferroni correction.

**Table S7:**
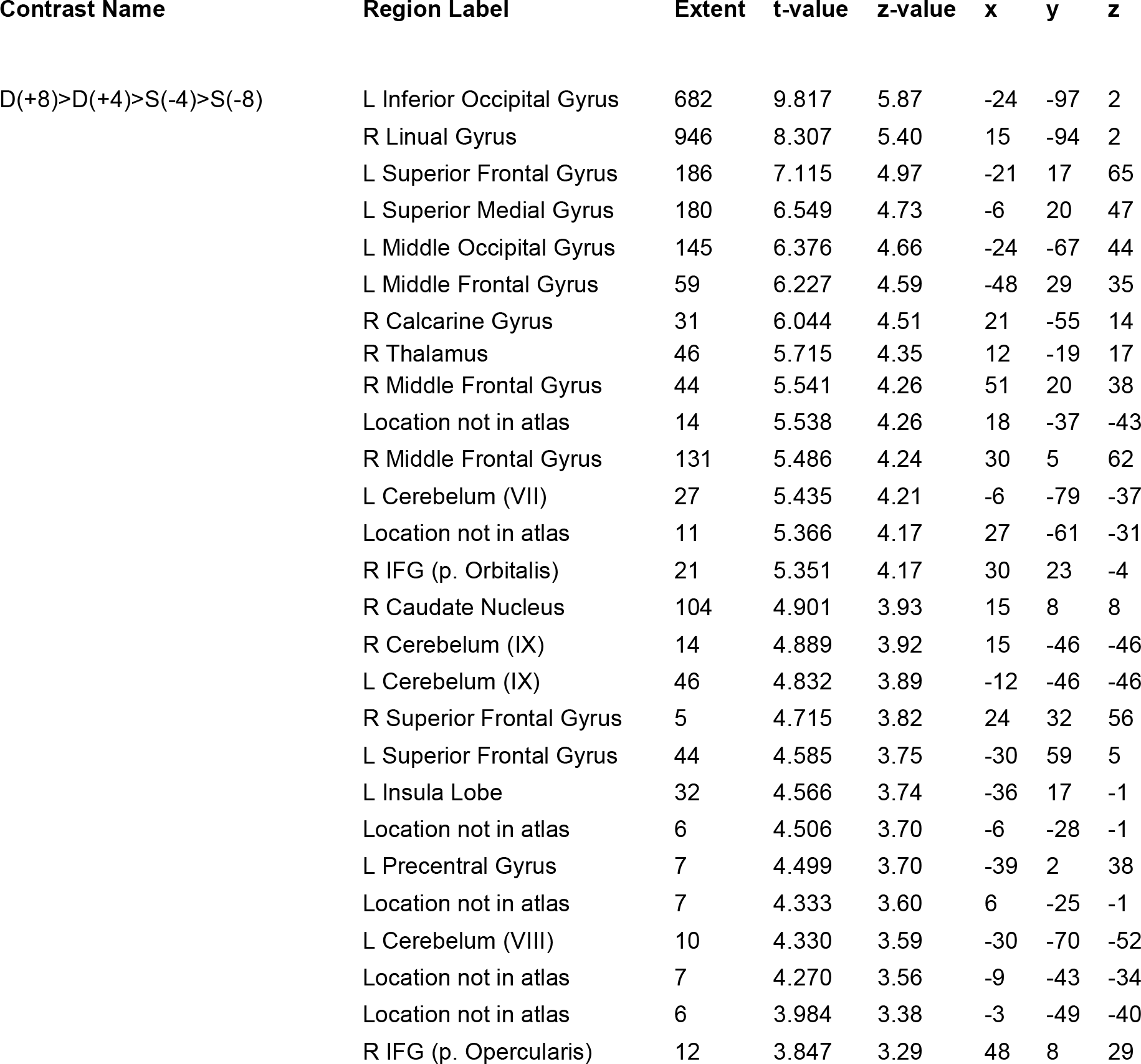
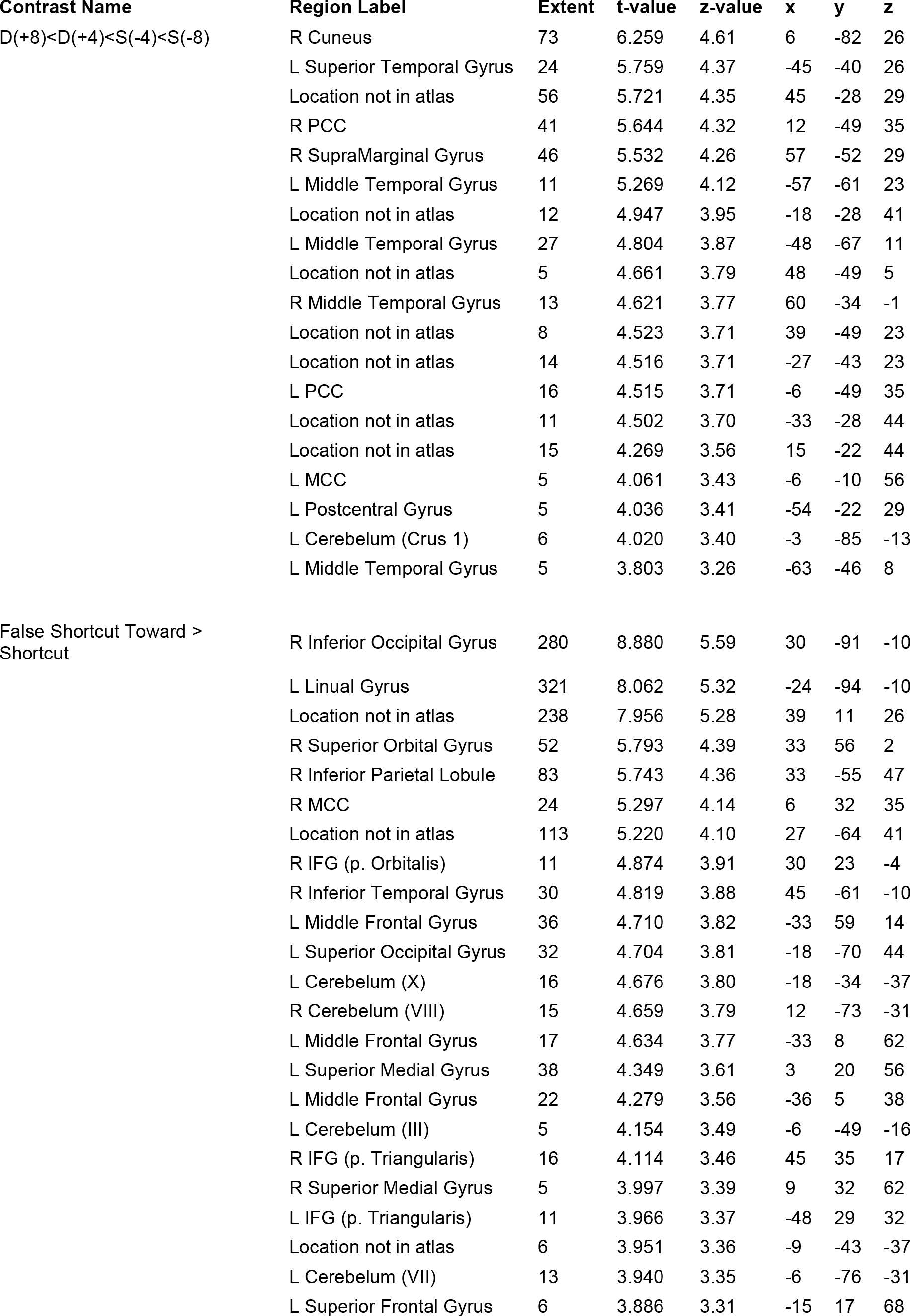
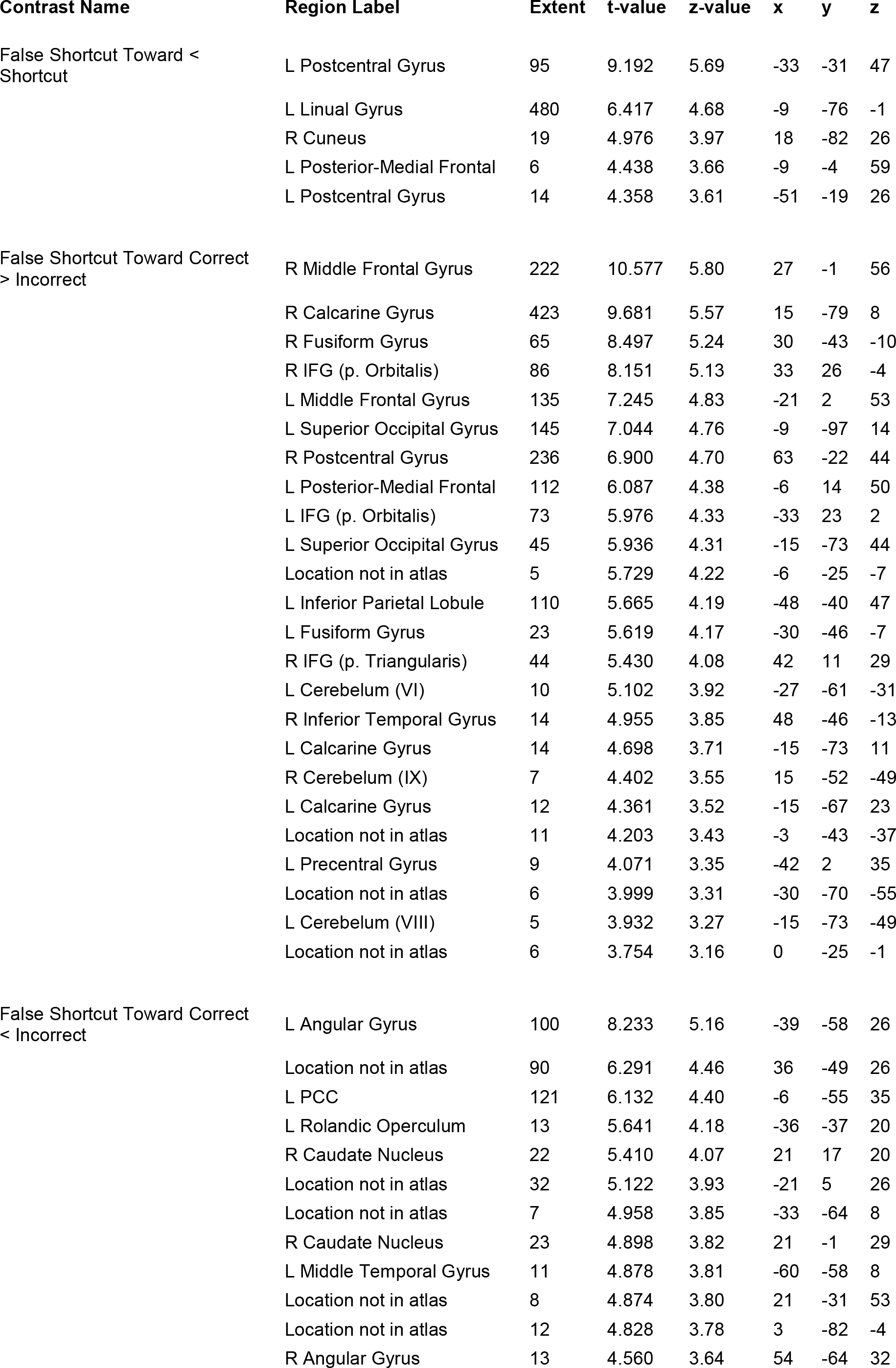
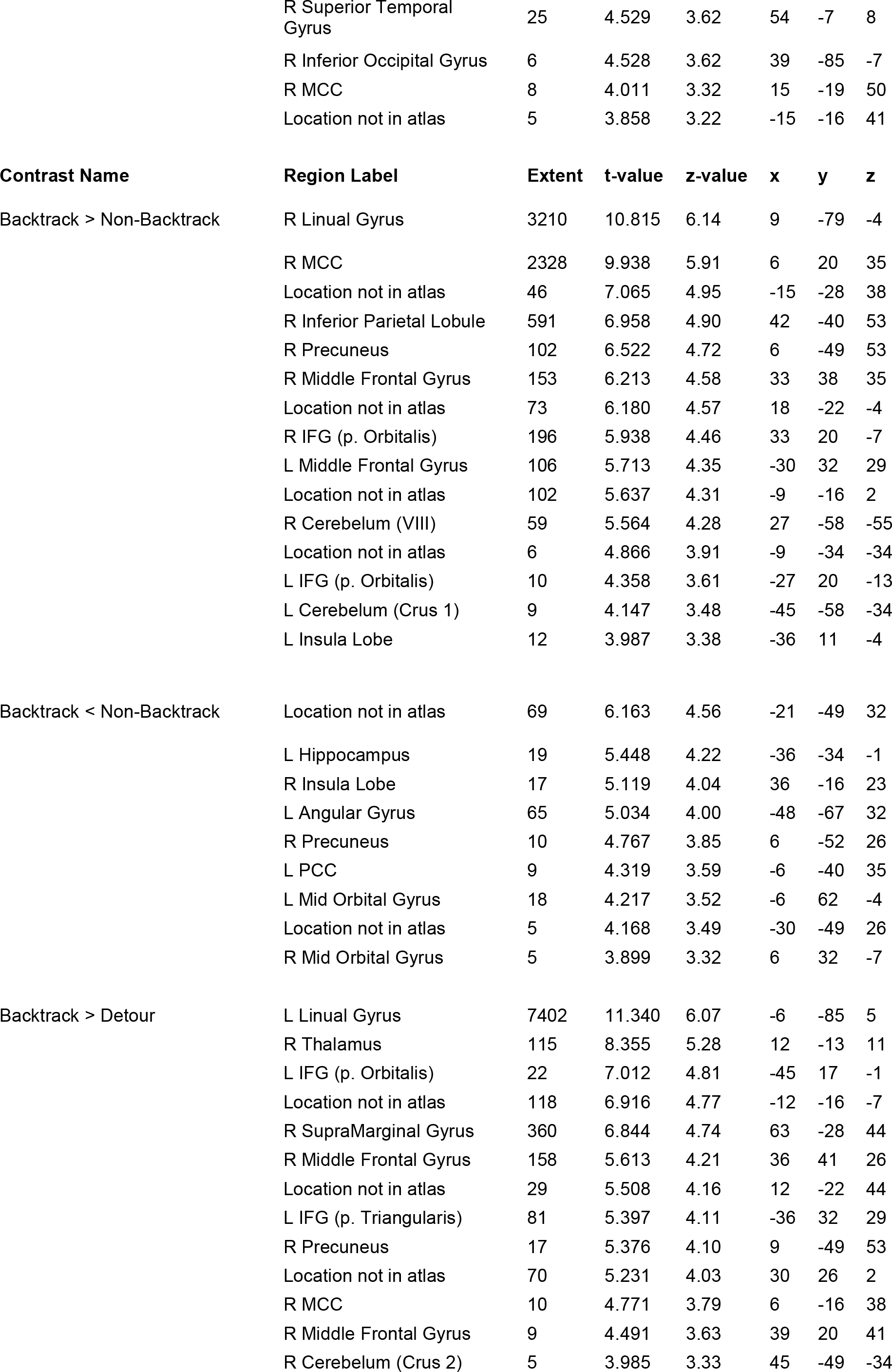
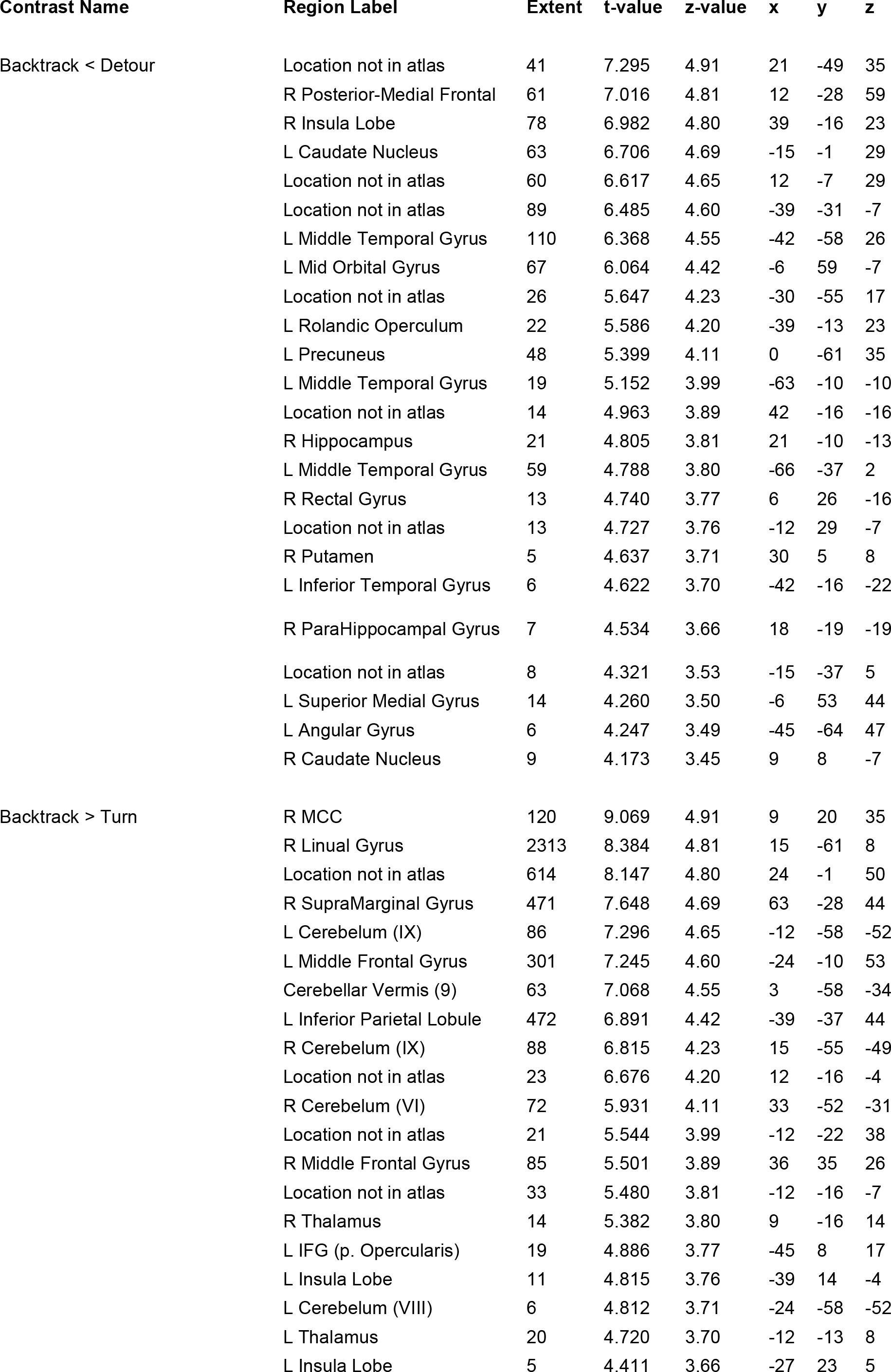
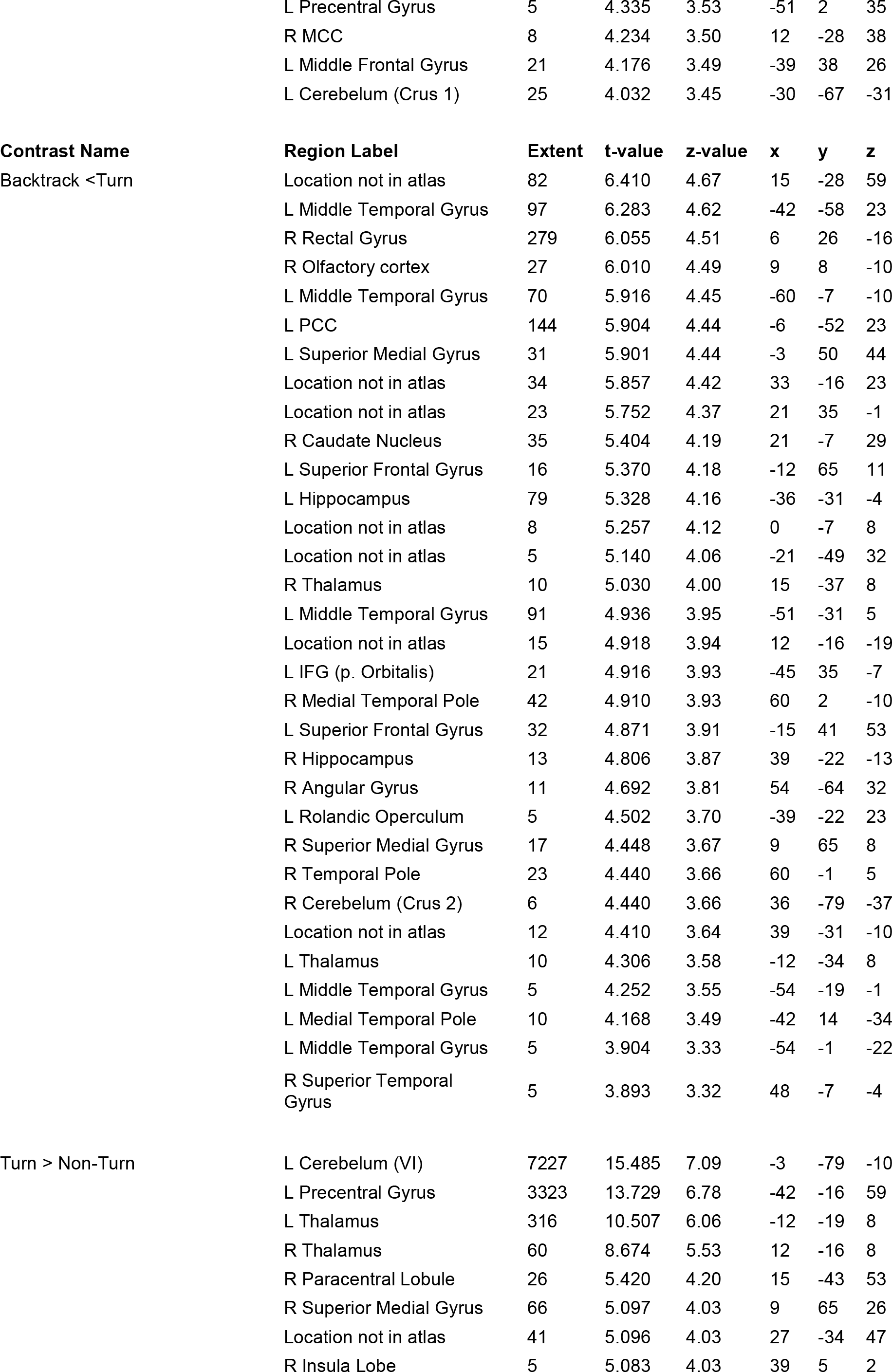
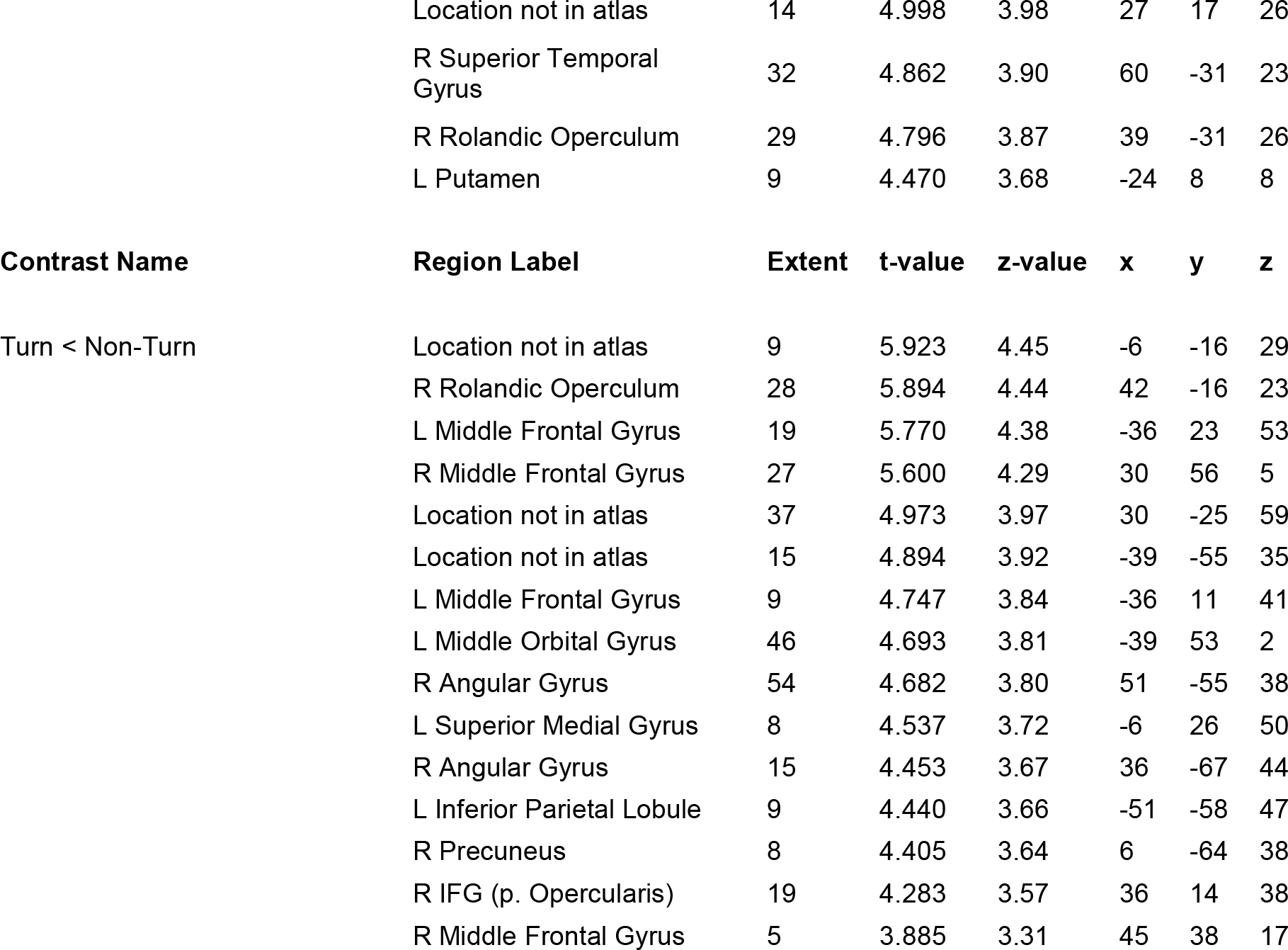
Summary of all fMRI activations. p<0.001 (uncorrected), min. 5 contiguous voxels Table shows all local maxima separated by more than 20 mm. Regions were automatically labeled using the AnatomyToolbox atlas using BSPMVIEW https://github.com/spunt/bspmview [https://zenodo.org/badge/latestdoi/21612/spunt/bspmview]

| Contrast Name | Region Label | Extent | t-value | z-value | x | y | z |
| --- | --- | --- | --- | --- | --- | --- | --- |
| D(+8)>D(+4)>S(-4)>S(-8) | L Inferior Occipital Gyrus | 682 | 9.817 | 5.87 | -24 | -97 | 2 |
|  | R Linual Gyrus | 946 | 8.307 | 5.40 | 15 | -94 | 2 |
|  | L Superior Frontal Gyrus | 186 | 7.115 | 4.97 | -21 | 17 | 65 |
|  | L Superior Medial Gyrus | 180 | 6.549 | 4.73 | -6 | 20 | 47 |
|  | L Middle Occipital Gyrus | 145 | 6.376 | 4.66 | -24 | -67 | 44 |
|  | L Middle Frontal Gyrus | 59 | 6.227 | 4.59 | -48 | 29 | 35 |
|  | R Calcarine Gyrus | 31 | 6.044 | 4.51 | 21 | -55 | 14 |
|  | R Thalamus | 46 | 5.715 | 4.35 | 12 | -19 | 17 |
|  | R Middle Frontal Gyrus | 44 | 5.541 | 4.26 | 51 | 20 | 38 |
|  | Location not in atlas | 14 | 5.538 | 4.26 | 18 | -37 | -43 |
|  | R Middle Frontal Gyrus | 131 | 5.486 | 4.24 | 30 | 5 | 62 |
|  | L Cerebelum (VII) | 27 | 5.435 | 4.21 | -6 | -79 | -37 |
|  | Location not in atlas | 11 | 5.366 | 4.17 | 27 | -61 | -31 |
|  | R IFG (p. Orbitalis) | 21 | 5.351 | 4.17 | 30 | 23 | -4 |
|  | R Caudate Nucleus | 104 | 4.901 | 3.93 | 15 | 8 | 8 |
|  | R Cerebelum (IX) | 14 | 4.889 | 3.92 | 15 | -46 | -46 |
|  | L Cerebelum (IX) | 46 | 4.832 | 3.89 | -12 | -46 | -46 |
|  | R Superior Frontal Gyrus | 5 | 4.715 | 3.82 | 24 | 32 | 56 |
|  | L Superior Frontal Gyrus | 44 | 4.585 | 3.75 | -30 | 59 | 5 |
|  | L Insula Lobe | 32 | 4.566 | 3.74 | -36 | 17 | -1 |
|  | Location not in atlas | 6 | 4.506 | 3.70 | -6 | -28 | -1 |
|  | L Precentral Gyrus | 7 | 4.499 | 3.70 | -39 | 2 | 38 |
|  | Location not in atlas | 7 | 4.333 | 3.60 | 6 | -25 | -1 |
|  | L Cerebelum (VIII) | 10 | 4.330 | 3.59 | -30 | -70 | -52 |
|  | Location not in atlas | 7 | 4.270 | 3.56 | -9 | -43 | -34 |
|  | Location not in atlas | 6 | 3.984 | 3.38 | -3 | -49 | -40 |
|  | R IFG (p. Opercularis) | 12 | 3.847 | 3.29 | 48 | 8 | 29 |

| **Contrast Name** | **Region Label** | **Extent** | **t-value** | **z-value** | **x** | **y** | **z** |
| --- | --- | --- | --- | --- | --- | --- | --- |
| D(+8)<D(+4)<S(-4)<S(-8) | R Cuneus | 73 | 6.259 | 4.61 | 6 | -82 | 26 |
|  | L Superior Temporal Gyrus | 24 | 5.759 | 4.37 | -45 | -40 | 26 |
|  | Location not in atlas | 56 | 5.721 | 4.35 | 45 | -28 | 29 |
|  | R PCC | 41 | 5.644 | 4.32 | 12 | -49 | 35 |
|  | R SupraMarginal Gyrus | 46 | 5.532 | 4.26 | 57 | -52 | 29 |
|  | L Middle Temporal Gyrus | 11 | 5.269 | 4.12 | -57 | -61 | 23 |
|  | Location not in atlas | 12 | 4.947 | 3.95 | -18 | -28 | 41 |
|  | L Middle Temporal Gyrus | 27 | 4.804 | 3.87 | -48 | -67 | 11 |
|  | Location not in atlas | 5 | 4.661 | 3.79 | 48 | -49 | 5 |
|  | R Middle Temporal Gyrus | 13 | 4.621 | 3.77 | 60 | -34 | -1 |
|  | Location not in atlas | 8 | 4.523 | 3.71 | 39 | -49 | 23 |
|  | Location not in atlas | 14 | 4.516 | 3.71 | -27 | -43 | 23 |
|  | L PCC | 16 | 4.515 | 3.71 | -6 | -49 | 35 |
|  | Location not in atlas | 11 | 4.502 | 3.70 | -33 | -28 | 44 |
|  | Location not in atlas | 15 | 4.269 | 3.56 | 15 | -22 | 44 |
|  | L MCC | 5 | 4.061 | 3.43 | -6 | -10 | 56 |
|  | L Postcentral Gyrus | 5 | 4.036 | 3.41 | -54 | -22 | 29 |
|  | L Cerebelum (Crus 1) | 6 | 4.020 | 3.40 | -3 | -85 | -13 |
|  | L Middle Temporal Gyrus | 5 | 3.803 | 3.26 | -63 | -46 | 8 |
| False Shortcut Toward > Shortcut | R Inferior Occipital Gyrus | 280 | 8.880 | 5.59 | 30 | -91 | -10 |
|  | L Linual Gyrus | 321 | 8.062 | 5.32 | -24 | -94 | -10 |
|  | Location not in atlas | 238 | 7.956 | 5.28 | 39 | 11 | 26 |
|  | R Superior Orbital Gyrus | 52 | 5.793 | 4.39 | 33 | 56 | 2 |
|  | R Inferior Parietal Lobule | 83 | 5.743 | 4.36 | 33 | -55 | 47 |
|  | R MCC | 24 | 5.297 | 4.14 | 6 | 32 | 35 |
|  | Location not in atlas | 113 | 5.220 | 4.10 | 27 | -64 | 41 |
|  | R IFG (p. Orbitalis) | 11 | 4.874 | 3.91 | 30 | 23 | -4 |
|  | R Inferior Temporal Gyrus | 30 | 4.819 | 3.88 | 45 | -61 | -10 |
|  | L Middle Frontal Gyrus | 36 | 4.710 | 3.82 | -33 | 59 | 14 |
|  | L Superior Occipital Gyrus | 32 | 4.704 | 3.81 | -18 | -70 | 44 |
|  | L Cerebelum (X) | 16 | 4.676 | 3.80 | -18 | -34 | -37 |
|  | R Cerebelum (VIII) | 15 | 4.659 | 3.79 | 12 | -73 | -31 |
|  | L Middle Frontal Gyrus | 17 | 4.634 | 3.77 | -33 | 8 | 62 |
|  | L Superior Medial Gyrus | 38 | 4.349 | 3.61 | 3 | 20 | 56 |
|  | L Middle Frontal Gyrus | 22 | 4.279 | 3.56 | -36 | 5 | 38 |
|  | L Cerebelum (III) | 5 | 4.154 | 3.49 | -6 | -49 | -16 |
|  | R IFG (p. Triangularis) | 16 | 4.114 | 3.46 | 45 | 35 | 17 |
|  | R Superior Medial Gyrus | 5 | 3.997 | 3.39 | 9 | 32 | 62 |
|  | L IFG (p. Triangularis) | 11 | 3.966 | 3.37 | -48 | 29 | 32 |
|  | Location not in atlas | 6 | 3.951 | 3.36 | -9 | -43 | -37 |
|  | L Cerebelum (VII) | 13 | 3.940 | 3.35 | -6 | -76 | -31 |
|  | L Superior Frontal Gyrus | 6 | 3.886 | 3.31 | -15 | 17 | 68 |
| False Shortcut Toward < Shortcut | L Postcentral Gyrus | 95 | 9.192 | 5.69 | -33 | -31 | 47 |
|  | L Linual Gyrus | 480 | 6.417 | 4.68 | -9 | -76 | -1 |
|  | R Cuneus | 19 | 4.976 | 3.97 | 18 | -82 | 26 |
|  | L Posterior-Medial Frontal | 6 | 4.438 | 3.66 | -9 | -4 | 59 |
|  | L Postcentral Gyrus | 14 | 4.358 | 3.61 | -51 | -19 | 26 |
| False Shortcut Toward Correct > Incorrect | R Middle Frontal Gyrus | 222 | 10.577 | 5.80 | 27 | -1 | 56 |
|  | R Calcarine Gyrus | 423 | 9.681 | 5.57 | 15 | -79 | 8 |
|  | R Fusiform Gyrus | 65 | 8.497 | 5.24 | 30 | -43 | -10 |
|  | R IFG (p. Orbitalis) | 86 | 8.151 | 5.13 | 33 | 26 | -4 |
|  | L Middle Frontal Gyrus | 135 | 7.245 | 4.83 | -21 | 2 | 53 |
|  | L Superior Occipital Gyrus | 145 | 7.044 | 4.76 | -9 | -97 | 14 |
|  | R Postcentral Gyrus | 236 | 6.900 | 4.70 | 63 | -22 | 44 |
|  | L Posterior-Medial Frontal | 112 | 6.087 | 4.38 | -6 | 14 | 50 |
|  | L IFG (p. Orbitalis) | 73 | 5.976 | 4.33 | -33 | 23 | 2 |
|  | L Superior Occipital Gyrus | 45 | 5.936 | 4.31 | -15 | -73 | 44 |
|  | Location not in atlas | 5 | 5.729 | 4.22 | -6 | -25 | -7 |
|  | L Inferior Parietal Lobule | 110 | 5.665 | 4.19 | -48 | -40 | 47 |
|  | L Fusiform Gyrus | 23 | 5.619 | 4.17 | -30 | -46 | -7 |
|  | R IFG (p. Triangularis) | 44 | 5.430 | 4.08 | 42 | 11 | 29 |
|  | L Cerebelum (VI) | 10 | 5.102 | 3.92 | -27 | -61 | -31 |
|  | R Inferior Temporal Gyrus | 14 | 4.955 | 3.85 | 48 | -46 | -13 |
|  | L Calcarine Gyrus | 14 | 4.698 | 3.71 | -15 | -73 | 11 |
|  | R Cerebelum (IX) | 7 | 4.402 | 3.55 | 15 | -52 | -49 |
|  | L Calcarine Gyrus | 12 | 4.361 | 3.52 | -15 | -67 | 23 |
|  | Location not in atlas | 11 | 4.203 | 3.43 | -3 | -43 | -37 |
|  | L Precentral Gyrus | 9 | 4.071 | 3.35 | -42 | 2 | 35 |
|  | Location not in atlas | 6 | 3.999 | 3.31 | -30 | -70 | -55 |
|  | L Cerebelum (VIII) | 5 | 3.932 | 3.27 | -15 | -73 | -49 |
|  | Location not in atlas | 6 | 3.754 | 3.16 | 0 | -25 | -1 |
| False Shortcut Toward Correct < Incorrect | L Angular Gyrus | 100 | 8.233 | 5.16 | -39 | -58 | 26 |
|  | Location not in atlas | 90 | 6.291 | 4.46 | 36 | -49 | 26 |
|  | L PCC | 121 | 6.132 | 4.40 | -6 | -55 | 35 |
|  | L Rolandic Operculum | 13 | 5.641 | 4.18 | -36 | -37 | 20 |
|  | R Caudate Nucleus | 22 | 5.410 | 4.07 | 21 | 17 | 20 |
|  | Location not in atlas | 32 | 5.122 | 3.93 | -21 | 5 | 26 |
|  | Location not in atlas | 7 | 4.958 | 3.85 | -33 | -64 | 8 |
|  | R Caudate Nucleus | 23 | 4.898 | 3.82 | 21 | -1 | 29 |
|  | L Middle Temporal Gyrus | 11 | 4.878 | 3.81 | -60 | -58 | 8 |
|  | Location not in atlas | 8 | 4.874 | 3.80 | 21 | -31 | 53 |
|  | Location not in atlas | 12 | 4.828 | 3.78 | 3 | -82 | -4 |
|  | R Angular Gyrus | 13 | 4.560 | 3.64 | 54 | -64 | 32 |

|  | R Superior Temporal Gyrus | 25 | 4.529 | 3.62 | 54 | -7 | 8 |
| --- | --- | --- | --- | --- | --- | --- | --- |
|  | R Inferior Occipital Gyrus | 6 | 4.528 | 3.62 | 39 | -85 | -7 |
|  | R MCC | 8 | 4.011 | 3.32 | 15 | -19 | 50 |
|  | Location not in atlas | 5 | 3.858 | 3.22 | -15 | -16 | 41 |
| Contrast Name | Region Label | Extent | t-value | z-value | x | y | z |
| Backtrack > Non-Backtrack | R Linual Gyrus | 3210 | 10.815 | 6.14 | 9 | -79 | -4 |
|  | R MCC | 2328 | 9.938 | 5.91 | 6 | 20 | 35 |
|  | Location not in atlas | 46 | 7.065 | 4.95 | -15 | -28 | 38 |
|  | R Inferior Parietal Lobule | 591 | 6.958 | 4.90 | 42 | -40 | 53 |
|  | R Precuneus | 102 | 6.522 | 4.72 | 6 | -49 | 53 |
|  | R Middle Frontal Gyrus | 153 | 6.213 | 4.58 | 33 | 38 | 35 |
|  | Location not in atlas | 73 | 6.180 | 4.57 | 18 | -22 | -4 |
|  | R IFG (p. Orbitalis) | 196 | 5.938 | 4.46 | 33 | 20 | -7 |
|  | L Middle Frontal Gyrus | 106 | 5.713 | 4.35 | -30 | 32 | 29 |
|  | Location not in atlas | 102 | 5.637 | 4.31 | -9 | -16 | 2 |
|  | R Cerebelum (VIII) | 59 | 5.564 | 4.28 | 27 | -58 | -55 |
|  | Location not in atlas | 6 | 4.866 | 3.91 | -9 | -34 | -34 |
|  | L IFG (p. Orbitalis) | 10 | 4.358 | 3.61 | -27 | 20 | -13 |
|  | L Cerebelum (Crus 1) | 9 | 4.147 | 3.48 | -45 | -58 | -34 |
|  | L Insula Lobe | 12 | 3.987 | 3.38 | -36 | 11 | -4 |
| Backtrack < Non-Backtrack | Location not in atlas | 69 | 6.163 | 4.56 | -21 | -49 | 32 |
|  | L Hippocampus | 19 | 5.448 | 4.22 | -36 | -34 | -1 |
|  | R Insula Lobe | 17 | 5.119 | 4.04 | 36 | -16 | 23 |
|  | L Angular Gyrus | 65 | 5.034 | 4.00 | -48 | -67 | 32 |
|  | R Precuneus | 10 | 4.767 | 3.85 | 6 | -52 | 26 |
|  | L PCC | 9 | 4.319 | 3.59 | -6 | -40 | 35 |
|  | L Mid Orbital Gyrus | 18 | 4.217 | 3.52 | -6 | 62 | -4 |
|  | Location not in atlas | 5 | 4.168 | 3.49 | -30 | -49 | 26 |
|  | R Mid Orbital Gyrus | 5 | 3.899 | 3.32 | 6 | 32 | -7 |
| Backtrack > Detour | L Linual Gyrus | 7402 | 11.340 | 6.07 | -6 | -85 | 5 |
|  | R Thalamus | 115 | 8.355 | 5.28 | 12 | -13 | 11 |
|  | L IFG (p. Orbitalis) | 22 | 7.012 | 4.81 | -45 | 17 | -1 |
|  | Location not in atlas | 118 | 6.916 | 4.77 | -12 | -16 | -7 |
|  | R SupraMarginal Gyrus | 360 | 6.844 | 4.74 | 63 | -28 | 44 |
|  | R Middle Frontal Gyrus | 158 | 5.613 | 4.21 | 36 | 41 | 26 |
|  | Location not in atlas | 29 | 5.508 | 4.16 | 12 | -22 | 44 |
|  | L IFG (p. Triangularis) | 81 | 5.397 | 4.11 | -36 | 32 | 29 |
|  | R Precuneus | 17 | 5.376 | 4.10 | 9 | -49 | 53 |
|  | Location not in atlas | 70 | 5.231 | 4.03 | 30 | 26 | 2 |
|  | R MCC | 10 | 4.771 | 3.79 | 6 | -16 | 38 |
|  | R Middle Frontal Gyrus | 9 | 4.491 | 3.63 | 39 | 20 | 41 |
|  | R Cerebelum (Crus 2) | 5 | 3.985 | 3.33 | 45 | -49 | -34 |
| Backtrack < Detour | Location not in atlas | 41 | 7.295 | 4.91 | 21 | -49 | 35 |
|  | R Posterior-Medial Frontal | 61 | 7.016 | 4.81 | 12 | -28 | 59 |
|  | R Insula Lobe | 78 | 6.982 | 4.80 | 39 | -16 | 23 |
|  | L Caudate Nucleus | 63 | 6.706 | 4.69 | -15 | -1 | 29 |
|  | Location not in atlas | 60 | 6.617 | 4.65 | 12 | -7 | 29 |
|  | Location not in atlas | 89 | 6.485 | 4.60 | -39 | -31 | -7 |
|  | L Middle Temporal Gyrus | 110 | 6.368 | 4.55 | -42 | -58 | 26 |
|  | L Mid Orbital Gyrus | 67 | 6.064 | 4.42 | -6 | 59 | -7 |
|  | Location not in atlas | 26 | 5.647 | 4.23 | -30 | -55 | 17 |
|  | L Rolandic Operculum | 22 | 5.586 | 4.20 | -39 | -13 | 23 |
|  | L Precuneus | 48 | 5.399 | 4.11 | 0 | -61 | 35 |
|  | L Middle Temporal Gyrus | 19 | 5.152 | 3.99 | -63 | -10 | -10 |
|  | Location not in atlas | 14 | 4.963 | 3.89 | 42 | -16 | -16 |
|  | R Hippocampus | 21 | 4.805 | 3.81 | 21 | -10 | -13 |
|  | L Middle Temporal Gyrus | 59 | 4.788 | 3.80 | -66 | -37 | 2 |
|  | R Rectal Gyrus | 13 | 4.740 | 3.77 | 6 | 26 | -16 |
|  | Location not in atlas | 13 | 4.727 | 3.76 | -12 | 29 | -7 |
|  | R Putamen | 5 | 4.637 | 3.71 | 30 | 5 | 8 |
|  | L Inferior Temporal Gyrus | 6 | 4.622 | 3.70 | -42 | -16 | -22 |
|  | R ParaHippocampal Gyrus | 7 | 4.534 | 3.66 | 18 | -19 | -19 |
|  | Location not in atlas | 8 | 4.321 | 3.53 | -15 | -37 | 5 |
|  | L Superior Medial Gyrus | 14 | 4.260 | 3.50 | -6 | 53 | 44 |
|  | L Angular Gyrus | 6 | 4.247 | 3.49 | -45 | -64 | 47 |
|  | R Caudate Nucleus | 9 | 4.173 | 3.45 | 9 | 8 | -7 |
| Backtrack > Turn | R MCC | 120 | 9.069 | 4.91 | 9 | 20 | 35 |
|  | R Linual Gyrus | 2313 | 8.384 | 4.81 | 15 | -61 | 8 |
|  | Location not in atlas | 614 | 8.147 | 4.80 | 24 | -1 | 50 |
|  | R SupraMarginal Gyrus | 471 | 7.648 | 4.69 | 63 | -28 | 44 |
|  | L Cerebelum (IX) | 86 | 7.296 | 4.65 | -12 | -58 | -52 |
|  | L Middle Frontal Gyrus | 301 | 7.245 | 4.60 | -24 | -10 | 53 |
|  | Cerebellar Vermis (9) | 63 | 7.068 | 4.55 | 3 | -58 | -34 |
|  | L Inferior Parietal Lobule | 472 | 6.891 | 4.42 | -39 | -37 | 44 |
|  | R Cerebelum (IX) | 88 | 6.815 | 4.23 | 15 | -55 | -49 |
|  | Location not in atlas | 23 | 6.676 | 4.20 | 12 | -16 | -4 |
|  | R Cerebelum (VI) | 72 | 5.931 | 4.11 | 33 | -52 | -31 |
|  | Location not in atlas | 21 | 5.544 | 3.99 | -12 | -22 | 38 |
|  | R Middle Frontal Gyrus | 85 | 5.501 | 3.89 | 36 | 35 | 26 |
|  | Location not in atlas | 33 | 5.480 | 3.81 | -12 | -16 | -7 |
|  | R Thalamus | 14 | 5.382 | 3.80 | 9 | -16 | 14 |
|  | L IFG (p. Opercularis) | 19 | 4.886 | 3.77 | -45 | 8 | 17 |
|  | L Insula Lobe | 11 | 4.815 | 3.76 | -39 | 14 | -4 |
|  | L Cerebelum (VIII) | 6 | 4.812 | 3.71 | -24 | -58 | -52 |
|  | L Thalamus | 20 | 4.720 | 3.70 | -12 | -13 | 8 |
|  | L Insula Lobe | 5 | 4.411 | 3.66 | -27 | 23 | 5 |
|  | L Precentral Gyrus | 5 | 4.335 | 3.53 | -51 | 2 | 35 |
|  | R MCC | 8 | 4.234 | 3.50 | 12 | -28 | 38 |
|  | L Middle Frontal Gyrus | 21 | 4.176 | 3.49 | -39 | 38 | 26 |
|  | L Cerebelum (Crus 1) | 25 | 4.032 | 3.45 | -30 | -67 | -31 |
| **Contrast Name** | **Region Label** | **Extent** | **t-value** | **z-value** | **x** | **y** | **z** |
| Backtrack <Turn | Location not in atlas | 82 | 6.410 | 4.67 | 15 | -28 | 59 |
|  | L Middle Temporal Gyrus | 97 | 6.283 | 4.62 | -42 | -58 | 23 |
|  | R Rectal Gyrus | 279 | 6.055 | 4.51 | 6 | 26 | -16 |
|  | R Olfactory cortex | 27 | 6.010 | 4.49 | 9 | 8 | -10 |
|  | L Middle Temporal Gyrus | 70 | 5.916 | 4.45 | -60 | -7 | -10 |
|  | L PCC | 144 | 5.904 | 4.44 | -6 | -52 | 23 |
|  | L Superior Medial Gyrus | 31 | 5.901 | 4.44 | -3 | 50 | 44 |
|  | Location not in atlas | 34 | 5.857 | 4.42 | 33 | -16 | 23 |
|  | Location not in atlas | 23 | 5.752 | 4.37 | 21 | 35 | -1 |
|  | R Caudate Nucleus | 35 | 5.404 | 4.19 | 21 | -7 | 29 |
|  | L Superior Frontal Gyrus | 16 | 5.370 | 4.18 | -12 | 65 | 11 |
|  | L Hippocampus | 79 | 5.328 | 4.16 | -36 | -31 | -4 |
|  | Location not in atlas | 8 | 5.257 | 4.12 | 0 | -7 | 8 |
|  | Location not in atlas | 5 | 5.140 | 4.06 | -21 | -49 | 32 |
|  | R Thalamus | 10 | 5.030 | 4.00 | 15 | -37 | 8 |
|  | L Middle Temporal Gyrus | 91 | 4.936 | 3.95 | -51 | -31 | 5 |
|  | Location not in atlas | 15 | 4.918 | 3.94 | 12 | -16 | -19 |
|  | L IFG (p. Orbitalis) | 21 | 4.916 | 3.93 | -45 | 35 | -7 |
|  | R Medial Temporal Pole | 42 | 4.910 | 3.93 | 60 | 2 | -10 |
|  | L Superior Frontal Gyrus | 32 | 4.871 | 3.91 | -15 | 41 | 53 |
|  | R Hippocampus | 13 | 4.806 | 3.87 | 39 | -22 | -13 |
|  | R Angular Gyrus | 11 | 4.692 | 3.81 | 54 | -64 | 32 |
|  | L Rolandic Operculum | 5 | 4.502 | 3.70 | -39 | -22 | 23 |
|  | R Superior Medial Gyrus | 17 | 4.448 | 3.67 | 9 | 65 | 8 |
|  | R Temporal Pole | 23 | 4.440 | 3.66 | 60 | -1 | 5 |
|  | R Cerebelum (Crus 2) | 6 | 4.440 | 3.66 | 36 | -79 | -37 |
|  | Location not in atlas | 12 | 4.410 | 3.64 | 39 | -31 | -10 |
|  | L Thalamus | 10 | 4.306 | 3.58 | -12 | -34 | 8 |
|  | L Middle Temporal Gyrus | 5 | 4.252 | 3.55 | -54 | -19 | -1 |
|  | L Medial Temporal Pole | 10 | 4.168 | 3.49 | -42 | 14 | -34 |
|  | L Middle Temporal Gyrus | 5 | 3.904 | 3.33 | -54 | -1 | -22 |
|  | R Superior Temporal Gyrus | 5 | 3.893 | 3.32 | 48 | -7 | -4 |
| Turn > Non-Turn | L Cerebelum (VI) | 7227 | 15.485 | 7.09 | -3 | -79 | -10 |
|  | L Precentral Gyrus | 3323 | 13.729 | 6.78 | -42 | -16 | 59 |
|  | L Thalamus | 316 | 10.507 | 6.06 | -12 | -19 | 8 |
|  | R Thalamus | 60 | 8.674 | 5.53 | 12 | -16 | 8 |
|  | R Paracentral Lobule | 26 | 5.420 | 4.20 | 15 | -43 | 53 |
|  | R Superior Medial Gyrus | 66 | 5.097 | 4.03 | 9 | 65 | 26 |
|  | Location not in atlas | 41 | 5.096 | 4.03 | 27 | -34 | 47 |
|  | R Insula Lobe | 5 | 5.083 | 4.03 | 39 | 5 | 2 |
|  | Location not in atlas | 14 | 4.998 | 3.98 | 27 | 17 | 26 |
|  | R Superior Temporal Gyrus | 32 | 4.862 | 3.90 | 60 | -31 | 23 |
|  | R Rolandic Operculum | 29 | 4.796 | 3.87 | 39 | -31 | 26 |
|  | L Putamen | 9 | 4.470 | 3.68 | -24 | 8 | 8 |
| **Contrast Name** | **Region Label** | **Extent** | **t-value** | **z-value** | **x** | **y** | **z** |
| Turn < Non-Turn | Location not in atlas | 9 | 5.923 | 4.45 | -6 | -16 | 29 |
|  | R Rolandic Operculum | 28 | 5.894 | 4.44 | 42 | -16 | 23 |
|  | L Middle Frontal Gyrus | 19 | 5.770 | 4.38 | -36 | 23 | 53 |
|  | R Middle Frontal Gyrus | 27 | 5.600 | 4.29 | 30 | 56 | 5 |
|  | Location not in atlas | 37 | 4.973 | 3.97 | 30 | -25 | 59 |
|  | Location not in atlas | 15 | 4.894 | 3.92 | -39 | -55 | 35 |
|  | L Middle Frontal Gyrus | 9 | 4.747 | 3.84 | -36 | 11 | 41 |
|  | L Middle Orbital Gyrus | 46 | 4.693 | 3.81 | -39 | 53 | 2 |
|  | R Angular Gyrus | 54 | 4.682 | 3.80 | 51 | -55 | 38 |
|  | L Superior Medial Gyrus | 8 | 4.537 | 3.72 | -6 | 26 | 50 |
|  | R Angular Gyrus | 15 | 4.453 | 3.67 | 36 | -67 | 44 |
|  | L Inferior Parietal Lobule | 9 | 4.440 | 3.66 | -51 | -58 | 47 |
|  | R Precuneus | 8 | 4.405 | 3.64 | 6 | -64 | 38 |
|  | R IFG (p. Opercularis) | 19 | 4.283 | 3.57 | 36 | 14 | 38 |
|  | R Middle Frontal Gyrus | 5 | 3.885 | 3.31 | 45 | 38 | 17 |

## Acknowledgements

We thank Mate Lengyel for advice on the experimental design. This work was supported by the Wellcome Trust (grant 094850/Z/10/Z) and James S. McDonnell Foundation to H.J.S, and the Gatsby Charitable Foundation (P.D.). The authors declare no competing financial interests. P.D. is currently on sabbatical at Uber AI Lab.

## Notes

Conflict of Interest: None.

## References

Alvernhe, A., Save, E., & Poucet, B. (2011). Local remapping of place cell firing in the Tolman detour task. European Journal of Neuroscience, 33(9), 1696–1705.

Alvernhe, A., Van Cauter, T., Save, E., & Poucet, B. (2008). Different CA1 and CA3 Representations of Novel Routes in a Shortcut Situation. Journal of Neuroscience, 28(29), 7324–7333.

Andrews-Hanna, J. R., Reidler, J. S., Sepulcre, J., Poulin, R., & Buckner, R. L. (2010). Functional-Anatomic Fractionation of the Brain’s Default Network. Neuron, 65(4), 550–562.

Balaguer, J., Spiers, H. J., Hassabis, D., & Summerfield, C. (2016). Neural Mechanisms of Hierarchical Planning in a Virtual Subway Network. Neuron, 90(4), 893–903.

Carter, C. S., Braver, T. S., Barch, D. M., Botvinick, M. M., Noll, D., & Cohen, J. D. (1998). Anterior cingulate cortex, error detection, and the online monitoring of performance. Science, 280(5364), 747–749.

Chapuis, N. (1987). Detour and Shortcut Abilities in Several Species of Mammals. In P. Ellen & C. Thinus-Blanc (Eds.), Cognitive Processes and Spatial Orientation in Animal and Man: Volume I Experimental Animal Psychology and Ethology (pp. 97–106). Dordrecht: Springer Netherlands.

Chapuis, N., Durup, M., & Thinus-Blanc, C. (1987). The role of exploratory experience in a shortcut task by golden hamsters (Mesocricetus auratus). Learning & Behavior, 15(2), 174–178.

Critchley, H. D., Wiens, S., Rotshtein, P., Öhman, A., & Dolan, R. J. (2004). Neural systems supporting interoceptive awareness. Nature Neuroscience, 7(2), 189–195.

Crittenden, B. M., Mitchell, D. J., & Duncan, J. (2015). Recruitment of the default mode network during a demanding act of executive control. ELife, 2015(4), 1–12.

Eichele, T., Debener, S., Calhoun, V. D., Specht, K., Engel, A. K., Hugdahl, K., … Ullsperger, M. (2008). Prediction of human errors by maladaptive changes in event-related brain networks. Proceedings of the National Academy of Sciences of the United States of America, 105(16), 6173–6178.

Ekstrom, A. D., Spiers, H. J., Bohbot, V. D., & Rosenbaum, S. R. (2018). HUMAN SPATIAL NAVIGATION. Princeton University Press.

Epstein, R. A., Patai, E. Z., Julian, J. B., & Spiers, H. J. (2017). The cognitive map in humans: Spatial navigation and beyond. Nature Neuroscience, 20(11), 1504–1513.

Fox, M. D., Snyder, A. Z., Barch, D. M., Gusnard, D. A., & Raichle, M. E. (2005). Transient BOLD responses at block transitions. NeuroImage, 28(4), 956–966.

Hassabis, D., & Maguire, E. A. (2007). Deconstructing episodic memory with construction. Trends in Cognitive Sciences, 11(7), 299–306.

Howard, L. R., Javadi, A.-H., Yu, Y., Mill, R. D., Morrison, L. C., Knight, R., … Spiers, H. J. (2014). The hippocampus and entorhinal cortex encode the path and Euclidean distances to goals during navigation. Current Biology, 24(12), 1331–40.

Iaria, G., Fox, C. J., Chen, J. K., Petrides, M., & Barton, J. J. S. (2008). Detection of unexpected events during spatial navigation in humans: Bottom-up attentional system and neural mechanisms. European Journal of Neuroscience, 27(4), 1017–1025.

Javadi, A.-H., Emo, B., Howard, L. R., Zisch, F. E., Yu, Y., Knight, R., … Spiers, H. J. (2017). Hippocampal and prefrontal processing of network topology to simulate the future. Nature Communications, 8, 14652.

Kaplan, R., King, J., Koster, R., Penny, W. D., Burgess, N., & Friston, K. J. (2017). The Neural Representation of Prospective Choice during Spatial Planning and Decisions. PLOS Biology, 15(1), e1002588.

Kolling, N., Wittmann, M. K., Behrens, T. E. J., Boorman, E. D., Mars, R. B., & Rushworth, M. F. S. (2016). Value, search, persistence and model updating in anterior cingulate cortex. Nature Neuroscience, 19(10), 1280–1285.

Kucyi, A., Hove, M. J., Esterman, M., Hutchison, R. M., & Valera, E. M. (2017). Dynamic Brain Network Correlates of Spontaneous Fluctuations in Attention. Cerebral Cortex (New York, N.Y.?: 1991), 27(3), 1831–1840.

Maguire, E. A., Burgess, N., Donnett, J. G., Frackowiak, R. S. J., Frith, C. D., & Keefe, J. O. (1998). Knowing Where and Getting There?: A Human Navigation Network. Science, 280(May), 921–924. Retrieved from http://www.ncbi.nlm.nih.gov/entrez/query.fcgi?cmd=Retrieve&db=PubMed&dopt=Citation&list_uids=9572740

Maldjian, J. A., Laurienti, P. J., Kraft, R. A., & Burdette, J. H. (2003). An automated method for neuroanatomic and cytoarchitectonic atlas-based interrogation of fMRI data sets. NeuroImage, 19(3), 1233–1239.

O’Doherty, J., Dayan, P., Schultz, J., Deichmann, R., Friston, K., & Dolan, R. J. (2004). Dissociable Roles of Ventral and Dorsal Striatum in Instrumental Conditioning. Science, 304(5669), 452–454.

O’Keefe, J., & Nadel, L. (1978). The Hippocampus as a Cognitive Map. Oxford University Press.

O’Reilly, J. X., Schuffelgen, U., Cuell, S. F., Behrens, T. E. J., Mars, R. B., & Rushworth, M. F. S. (2013). Dissociable effects of surprise and model update in parietal and anterior cingulate cortex. Proceedings of the National Academy of Sciences, 110(38), E3660–E3669.

Ólafsdóttir, H. F., Barry, C., Saleem, A. B., Hassabis, D., & Spiers, H. J. (2015). Hippocampal place cells construct reward related sequences through unexplored space. ELife, 4, 1–17.

Patai, E. Z., Javadi, A.-H., Ozubko, J. D., Ji, S., O’Callaghan, A., Robin, J., … Spiers, H. J. (2017). Long-term consolidation switches goal proximity coding from hippocampus to retrosplenial cortex. BioRxiv. Retrieved from http://www.biorxiv.org/content/early/2017/07/25/167882

Pfeiffer, B. E., & Foster, D. J. (2013). Hippocampal place-cell sequences depict future paths to remembered goals. Nature, 497(7447), 1–8.

Poucet, B., Thinus-Blanc, C., & Chapuis, N. (1983). Route planning in cats, in relation to the visibility of the goal. Animal Behaviour, 31(2), 594–599.

Rauchs, G., Orban, P., Balteau, E., Schmidt, C., Degueldre, C., Luxen, A., … Peigneux, P. (2008). Partially segregated neural networks for spatial and contextual memory in virtual navigation. Hippocampus, 18(5), 503–518.

Rosenbaum, S. R., Ziegler, M., Winocur, G., Grady, C. L., & Moscovitch, M. (2004). “I have often walked down this street before”: fMRI studies on the hippocampus and other structures during mental navigation of an old environment. Hippocampus, 14(7), 826–835.

Seeley, W. W., Menon, V., Schatzberg, A. F., Keller, J., Glover, G. H., Kenna, H., … Greicius, M. D. (2007). Dissociable Intrinsic Connectivity Networks for Salience Processing and Executive Control. Journal of Neuroscience, 27(9), 2349–2356.

Shallice, T. (1982). Specific impairments of planning. Philosophical Transactions of the Royal Society, 298.

Simon, D. A., & Daw, N. D. (2011). Neural correlates of forward planning in a spatial decision task in humans. The Journal of Neuroscience, 31(14), 5526–5539.

Smith, V., Mitchell, D. J., & Duncan, J. (2018). Role of the Default Mode Network in Cognitive Transitions. Cerebral Cortex, (August).

Spiers, H. J. (2008). Keeping the goal in mind: Prefrontal contributions to spatial navigation. Neuropsychologia, 46(7), 2106–2108.

Spiers, H. J., & Barry, C. (2015). Neural systems supporting navigation. Current Opinion in Behavioral Sciences, 1, 47–55.

Spiers, H. J., & Gilbert, S. J. (2015). Solving the detour problem in navigation: a model of prefrontal and hippocampal interactions. Frontiers in Human Neuroscience, 9(March), 1–15.

Spiers, H. J., & Maguire, E. A. (2006). Thoughts, behaviour, and brain dynamics during navigation in the real world. NeuroImage, 31(4), 1826–1840.

Sridharan, D., Levitin, D. J., & Menon, V. (2008). A critical role for the right fronto-insular cortex in switching between central-executive and default-mode networks. Proceedings of the National Academy of Sciences, 105(34), 12569–12574.

Tolman, E. C. (1948). Cognitive maps in rats and men. Psychological Review, 55(4), 189–208.

Tolman, E. C., & Honzik, C. H. (1930). Introduction and removal of reward and maze performance in rats. Berkeley, Calif.: University of California Press.

Tzourio-Mazoyer, N., Landeau, B., Papathanassiou, D., Crivello, F., Etard, O., Delcroix, N., … Joliot, M. (2002). Automated Anatomical Labeling of Activations in SPM Using a Macroscopic Anatomical Parcellation of the MNI MRI Single-Subject Brain. NeuroImage, 15(1), 273–289.

Wang, C., Ulbert, I., Schomer, D. L., Marinkovic, K., & Halgren, E. (2005). Responses of Human Anterior Cingulate Cortex Microdomains to Error Detection, Conflict Monitoring, Stimulus-Response Mapping, Familiarity, and Orienting. Journal of Neuroscience, 25(3), 604–613.

Winocur, G., Moscovitch, M., Rosenbaum, S. R., & Sekeres, M. (2010). An investigation of the effects of hippocampal lesions in rats on pre- and postoperatively acquired spatial memory in a complex environment. Hippocampus, 20(12), 1350–1365.

Wystrach, A., Schwarz, S., Baniel, A., & Cheng, K. (2013). Backtracking behaviour in lost ants: an additional strategy in their navigational toolkit. Proceedings. Biological Sciences / The Royal Society, 280(1769), 20131677.

Xu, J., Evensmoen, H. R., Lehn, H., Pintzka, C. W. S., & Håberg, A. K. (2010). Persistent posterior and transient anterior medial temporal lobe activity during navigation. NeuroImage, 52(4), 1654–1666.

Yoshida, W., & Ishii, S. (2006). Resolution of Uncertainty in Prefrontal Cortex. Neuron, 50(5), 781–789.

